# Dietary tryptophan and genetic susceptibility expand gut microbiota that promote systemic autoimmune activation

**DOI:** 10.1101/2024.01.16.575942

**Authors:** Longhuan Ma, Yong Ge, Josephine Brown, Seung-Chul Choi, Ahmed Elshikha, Nathalie Kanda, Morgan Terrell, Natalie Six, Abigail Garcia, Mansour Mohamadzadeh, Gregg Silverman, Laurence Morel

## Abstract

Tryptophan modulates disease activity and the composition of microbiota in the B6.*Sle1.Sle2.Sle3* (TC) mouse model of lupus. To directly test the effect of tryptophan on the gut microbiome, we transplanted fecal samples from TC and B6 control mice into germ-free or antibiotic-treated non-autoimmune B6 mice that were fed with a high or low tryptophan diet. The recipient mice with TC microbiota and high tryptophan diet had higher levels of immune activation, autoantibody production and intestinal inflammation. A bloom of *Ruminococcus gnavus (Rg),* a bacterium associated with disease flares in lupus patients, only emerged in the recipients of TC microbiota fed with high tryptophan. *Rg* depletion in TC mice decreased autoantibody production and increased the frequency of regulatory T cells. Conversely, TC mice colonized with *Rg* showed higher autoimmune activation. Overall, these results suggest that the interplay of genetic and tryptophan can influence the pathogenesis of lupus through the gut microbiota.

## Introduction

Systemic lupus erythematosus (SLE) is an autoimmune disease characterized by the generation of autoantibodies directed against nucleic acid / protein complexes responsible for tissue damage in multiple organs^1^. The etiology of SLE involves a combination of genetic, epigenetic and environmental factors^2,3^ ^4^. The gut microbiota is one type of environmental factors that has been closely associated to SLE^5^. Decreased richness and diversity of gut microbiota along with variations in specific microbial taxa have been reported in SLE patients and animal models of the disease^6^. Individual bacterium, such as *Lactobacillus reuteri*^7^ and *Enteroccocus gallinarium*^8^ have been identified as pathobionts contributing to autoimmune activation in mouse models of SLE. The abundance of fecal *Ruminococcus (blautia) gnavus* (*Rg*) was positively correlated with disease severity in lupus nephritis patients^9^. Moreover, longitudinal studies showed that bacterial blooms occurred during disease flares, with *Rg* being the most consistently expanded taxa^10^. *Rg* produces a highly immunogenic lipoglycan (LP), and high titers of anti-LP IgG2 also correlated with disease activity in these patients^10^. However, the cause of gut dysbiosis in general and the expansion of specific bacteria such as *Rg* in SLE patients are unknown. Gut microbial dysbiosis could be a cause or a consequence of an abnormal host immune state, since microbes can modulate the immune system, and immune activation can in turn impact the microbiome architecture^11^. It is also recognized that genetic susceptibility contributes to autoimmune activation in lupus^12^, and therefore it could indirectly shape the gut microbiome.

Gut microbes can impact the host immune state by directly interacting with immune cells as whole bacteria or with their components, including the metabolites they produce. In addition, an unbalanced gut microbiota can also skew the metabolism of consumed food or supplements, which may change the profile of immune modulatory components^13^. We have reported a gut microbial dysbiosis and a skewed tryptophan metabolism that is associated with autoimmune activation in B6.*Sle1.Sle2*.*Sle3* triple congenic (TC) lupus-prone mice as compared to their congenic healthy B6 controls^14^. TC mice presented a distinct profile of tryptophan metabolites in their serum, feces as well as CD4^+^ T cells as compared to congenic B6 control mice ^14,15^. Similar levels of endogenous tryptophan metabolic enzymes are expressed by TC and B6 mice, suggesting that the skewed tryptophan catabolism in lupus-prone mice may largely be of bacterial origin^15^. Tryptophan and its bacterial-produced metabolite, tryptamine, promoted mTOR activation and glycolysis in isolated CD4^+^ T cells from TC mice, but from not B6 mice^15^, suggesting a direct immune-regulatory effect that depends on genetic susceptibility.

An altered tryptophan metabolism has been reported in the serum, immune cells, and feces of SLE patients in correlation with disease activity^16–22^. High dietary tryptophan exacerbated lupus phenotypes in TC mice, while low dietary tryptophan delayed the progression of disease, including an expansion of regulatory FOXP3^+^ CD4^+^ T (Treg) cells^14^. Variations in dietary tryptophan also induced specific alterations in the gut microbiome of TC relative to B6 mice, with exposure to high tryptophan expanding some of the taxa that differ between the two strains on standard chow. The disease modulatory function of tryptophan could be mediated through a skewed bacterial metabolism or through direct interactions with the activated lupus immune cells. To address this question, we excluded the direct contribution of lupus genetic susceptibility on the immune system and potential differences in intrinsic tryptophan metabolisms. We performed fecal microbiota transfers (FMT) from TC or B6 mice into antibiotic-treated B6 recipient mice, which were fed a high or low dietary tryptophan diet. The combination of TC microbiota and high tryptophan diet induced autoimmune responses and gut inflammation in B6 recipients. These observations suggest that both high dietary tryptophan and components of the microbiota that develop in TC mice contribute to autoimmune activation. Distinct fecal metabolomic profiles were observed in the recipient mice based on microbiota origins and dietary tryptophan contents. Moreover, using germ-free B6 recipients, we showed that high tryptophan enriched unique microbial taxa in the gut microbiota of TC or B6 origin, with *Rg* being enriched only in recipients of TC microbiota. Coincidently, we observed higher *Rg* levels in the feces of TC mice compared to age-matched B6 mice, and *Rg* abundance was positively correlated with serum IL-6 and IFN-γ, two cytokines found at high levels in SLE ^23,24^. Furthermore, *Rg* depletion in TC mice decreased the production of autoantibodies while increasing the frequency of Treg cells, while *Rg* monocolonization of TC mice induced autoimmune activation and autoantibody production. Finally, stimulation of immune cells in the presence of *Rg* lysate induced Treg cell apoptosis. These observations suggest that the combination of altered tryptophan metabolism and activated adaptive immunity enriches the *Rg* population in the gut of lupus-prone mice, which promotes autoimmunity, at least in part by targeting Treg cells. Considering the positive correlation between the abundance of fecal *Rg* and disease activity in SLE patients^10^, the results suggest a model in which genetic susceptibility drives the development of an inflammatory immune system that shapes the microbiome towards an altered tryptophan catabolism. This in turn favors the expansion of *Rg* populations, which independently activate autoimmune responses.

## Results

### The combination of gut microbiota from lupus-prone mice and high dietary tryptophan promoted autoimmune activation

To eliminate the direct effect of lupus-suceptibility genes on immune activation or intrinsic tryptophan metabolism, we performed fecal microbiota transfers (FMT) from TC mice with high anti-dsDNA IgG level or age-matched B6 controls into B6 recipients that were previoulsy treated with broad-spectrum antibiotics, which greatly reduced their fecal bacterial load (**Figure S1A**). The recipent mice were fed with high (1.19%) or low tryptophan (∼ 0.03%) synthetic diets for 4 weeks starting one week before FMT. This experimental design resulted in 4 groups: high tryptophan with TC microbiota (TCmbTrp^hi^), low tryptophan with TC microbiota (TCmbTrp^lo^), high tryptophan with B6 microbiota (B6mbTrp^hi^), and low tryptophan with B6 microbiota (B6mbTrp^lo^) (**Figure S1B**). We have previously reported that low dietary tryptophan induced a rapid body weight loss in TC mice ^14^. Here, only TCmbTrp^lo^ mice suffered weight loss (**Figure 1A**), suggesting that this process is driven by TC microbiota consuming more rapidly the low amount of tryptophan. TCmbTrp^hi^ mice showed a higher spleen weight to bodyweight ratio than B6mbTrp^hi^ mice, suggesting a higer systemic inflammation (**Figure 1B**). TCmbTrp^hi^ mice produced more serum total IgA and anti-dsDNA IgA than either B6mbTrp^hi^ or B6mbTrp^lo^ mice, with intermediate values in TCmbTrp^lo^ mice (**Figure 1C**), and TCmbTrp^hi^ mice were the only group to produce high level of cecal anti-dsDNA IgA (**Figure 1D**). Total IgA was also the most abundant in cecal content of TC microbiota recipients, and it was decreased by low tryptophan (**Figure 1E**). No difference were detected between groups for serum anti-dsDNA IgG by ELISA. However, the highly sensitive and specific *Crithidia luciliae* indirect immunofluorescence assay detected a higher level of anti-dsDNA IgG in the serum of TCmbTrp^hi^ mice compared to all the other groups (**Figure 1F**). In addition, the serum anti-dsDNA IgA / IgM ratio was higher in TCmbTrp^hi^ mice than in the three other groups, and there was a similar trend for the anti-dsDNA IgG / IgM ratio (**Figure S1C-D**). Since these two ratios have been correlated to lupus nephritis (LN), and considering that T cells are a major pathogenic contributor to LN^25^, we assessed T cell inflitration in the kidneys. A higher number of CD3^+^ T cells infiltrated the kidneys of TCmbTrp^hi^ mice than in the three other groups (**Figure 1G**).

**Figure 1.**
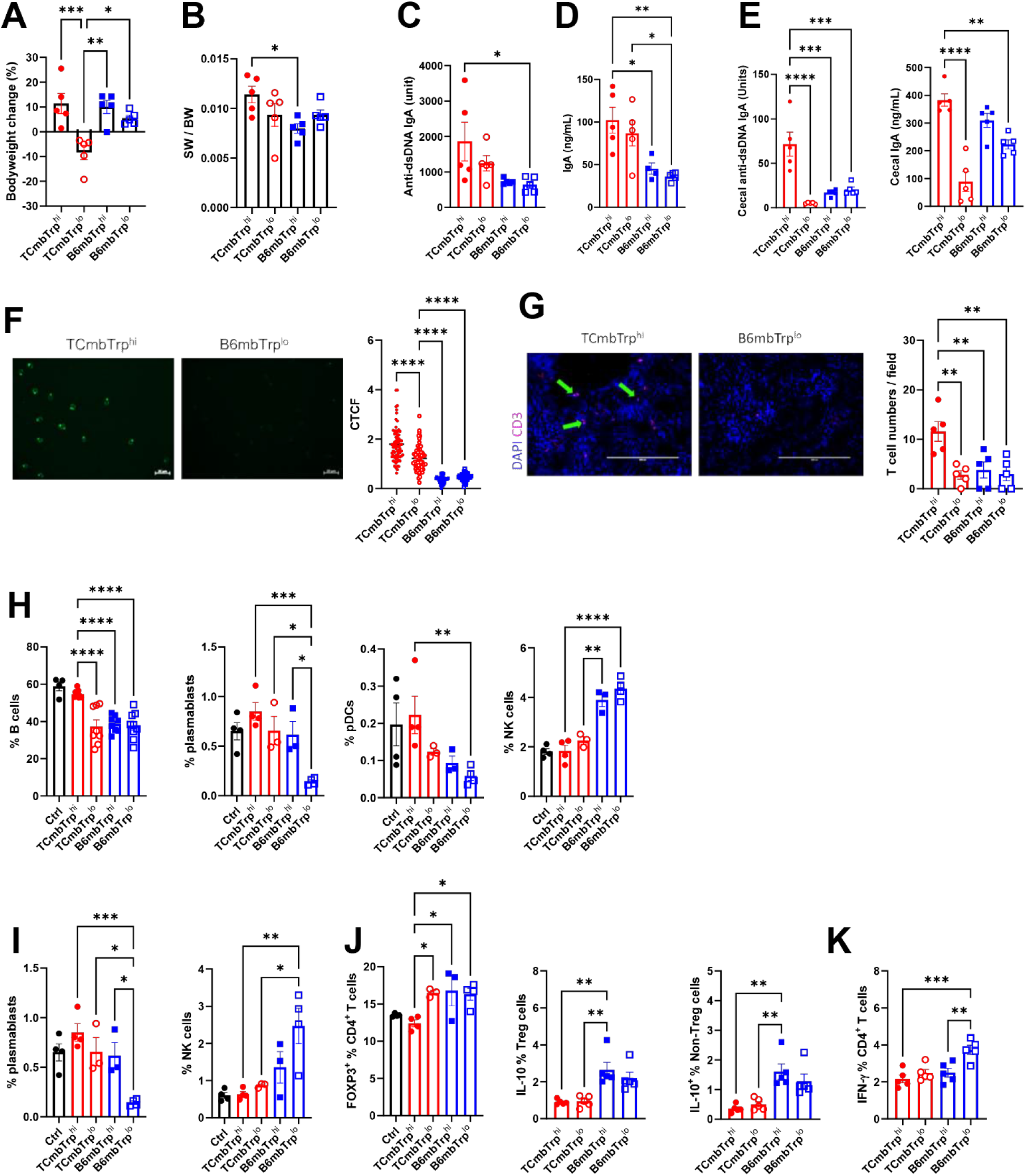
The combination of gut microbiota from lupus-prone mice and high dietary tryptophan promotes autoimmune activation in non-autoimmune FMT recipients. (A) Terminal to initial bodyweight changes. (B) Spleen weight (SW) / bodyweight (BW) ratio. Terminal serum anti-dsDNA IgA (C) and total IgA (D). (E) Terminal fecal anti-dsDNA IgA and cecal total IgA. (F) Representative images (Scale bar: 20 µm) and quantitation of a *Crithidia luciliae* assay on terminal serum. Corrected total cell fluorescence (CTCF) was calculated on all visible IgG^+^ cells in each slide. (G) Representative images (scale bars: 200 µm) and quantitation of CD3^+^ cell infiltrates in the kidney. All CD3^+^ cells were counted in the 20X field. (H) Frequency of B cells, plasmablasts, NK cells and pDCs in the spleen. (I) Frequency of plasmablasts and NK cells in the mLN. (J) Frequency of mLN Treg cells in CD4^+^ T cells, IL-10^+^ cells in Treg and non-Treg CD4^+^ T cells. (K) Frequency of IFN-γ^+^ CD4^+^ T cells in the mLN. N = 3 – 9 FMT recipients per group. Control (Ctrl) mice (black symbols in H – J) were untouched age-matched B6 mice. Mean + SEM compared with 1-way ANOVA with multiple-comparison tests. *: P < 0.05; **: P < 0.01; ***: P < 0.001; ****: P < 0.001.

In the spleen, a higher frequency of B cells, plasmablasts and plasmacytoid DCs (pDCs), three immune cell types involved in SLE, was found in TCmbTrp^hi^ mice (**Figure 1H and Figure S2** for gating strategy). On the other hand, the B6 microbiota, regardless of tryptophan, increased the frequency of splenic NK cells (**Figure 1H**). An increased frequency of plasmablasts and a decreased frequency of NK cells was confirmed in the mesenteri lymph node (mLN) of TCmbTrp^hi^ mice (**Figure 1I**). The role of NK cells in autoimmunity is complex, but their reduced frfequency and protective function have been demonstrated in a mouse model of lupus^26^. The distribution of the major CD4^+^ T cell effector subsets in the spleen was largely similar between the four recipient groups . However, differences were observed in mLN, where immune cells were most likely directly impacted by the transferred microbiota and dietary tryptophan. A lower frequency of FOXP3^+^ Treg cells was observed in TCmbTrp^hi^ mice (**Figure 1J**). In addition, a lower frequency of IL-10-producing Treg and non-Treg CD4^+^ T cells was observed in groups of mice that received TC microbiota regardless of dietary tryptophan (**Figure 1J**). Surprisingly, a higher frequency of IFN-γ^+^ CD4^+^ T cells was also detected in the mLN of B6mbTrp^lo^ mice (**Figure 1K**). Although there is strong evidence that IFN-γ is pathogenenic in SLE^27^, an anti-inflammatory role has also been reported for this cytokine^28,29^, which may be involved in the FMT experimental setting. Overall, these results suggest that the origin of the microbiota and dietary tryptophan modulate the immune system, with the combination of gut microbiota originating in autoimmune TC mice and high dietary tryptophan promoting autoimmune phenotypes on a non-autoimmune genetic background.

### The combination of gut microbiota from lupus-prone mice and high dietary tryptophan promoted intestinal inflammation

SLE has been associated with a disrupted intestinal barrier in patients and several mouse models of the disease^6,30^. However, lupus-prone TC mice do not present a loss of intestinal barrier integrity^14^ unless they are treated with a TLR7 agonist^31^. Here, we assessed gut permeability in the four groups of FMT recipient mice by measuring orally administrated 4-kDa FITC-dextran in the serum. Results suggested that the transfer of TC gut microbiota increased gut permeability in this model independently from dietary tryptophan (**Figure 2A**). Elevated serum zonulin is a marker of impaired tight junctions and gut barrier integrity ^32^. The lowest level of serum zonulin (**Figure 2B**) and the highest expression of the tight junction protein, Claudin-1, found in the small intestine (**Figure 2C**) of B6mbTrp^lo^ mice suggested that both the TC origin of the microbiota and high dietary tryptophan enhanced gut permeability. Additionally, TCmbTrp^hi^ mice developed the highest number of CD45^+^ leukocytes and immune cell foci in the colon (**Figure 2D**). The identity and frequency of the immune cells in the colon lamina propria was analyzed by flow cytometry in the four groups of FMT recipients (**Figure S3** for gating strategy). B6mbTrp^lo^ mice presented the lowest frequency of B cells (**Figure 2E**), and the frequency of CD4^+^ T cells was increased by low dietary tryptophan regardless of the microbiota origin (**Figure 2F**). Among the CD4^+^ T cells, the recipients of TC microbiota presented a lower frequency of IL-17a-producing cells and CD3e^-^ NK1.1^+^ NK cells regardless of tryptophan(**Figure 2G-H**). These results are consistent with the recipients of TC microbiota presenting a lower gut barrier integrity, since IL-17a^+^ T cells^33^ and NK cells^34^ play a protective role in this process. Finally, the frequency of conventional DCs (cDCs) was the lowest in TCmbTrp^hi^ mice and the highest in B6mbTrp^lo^ mice (**Figure 2I**), while the frequency of inflammatory pDCs (ipDCs) was higher in the recipients of B6 microbiota (**Figure 2J**). These results are consistent with the frequency of NK cells, considering the positive feedback loop between NK cells, DCs and pDCs^35^. Consistent with the high production of IgA and anti-dsDNA IgA, a large number of IgA^+^ cells were detected in the colon of recipients of TC microbiota that was enhanced by dietary tryptophan (**Figure 2K**), suggesting an increased local IgA production induced by gut inflammation^36^. Taken together, these results showed that TC-derived microbiota drives gut inflammation and alters barrier integrity, with some modulation from dietary tryptophan.

**Figure 2.**
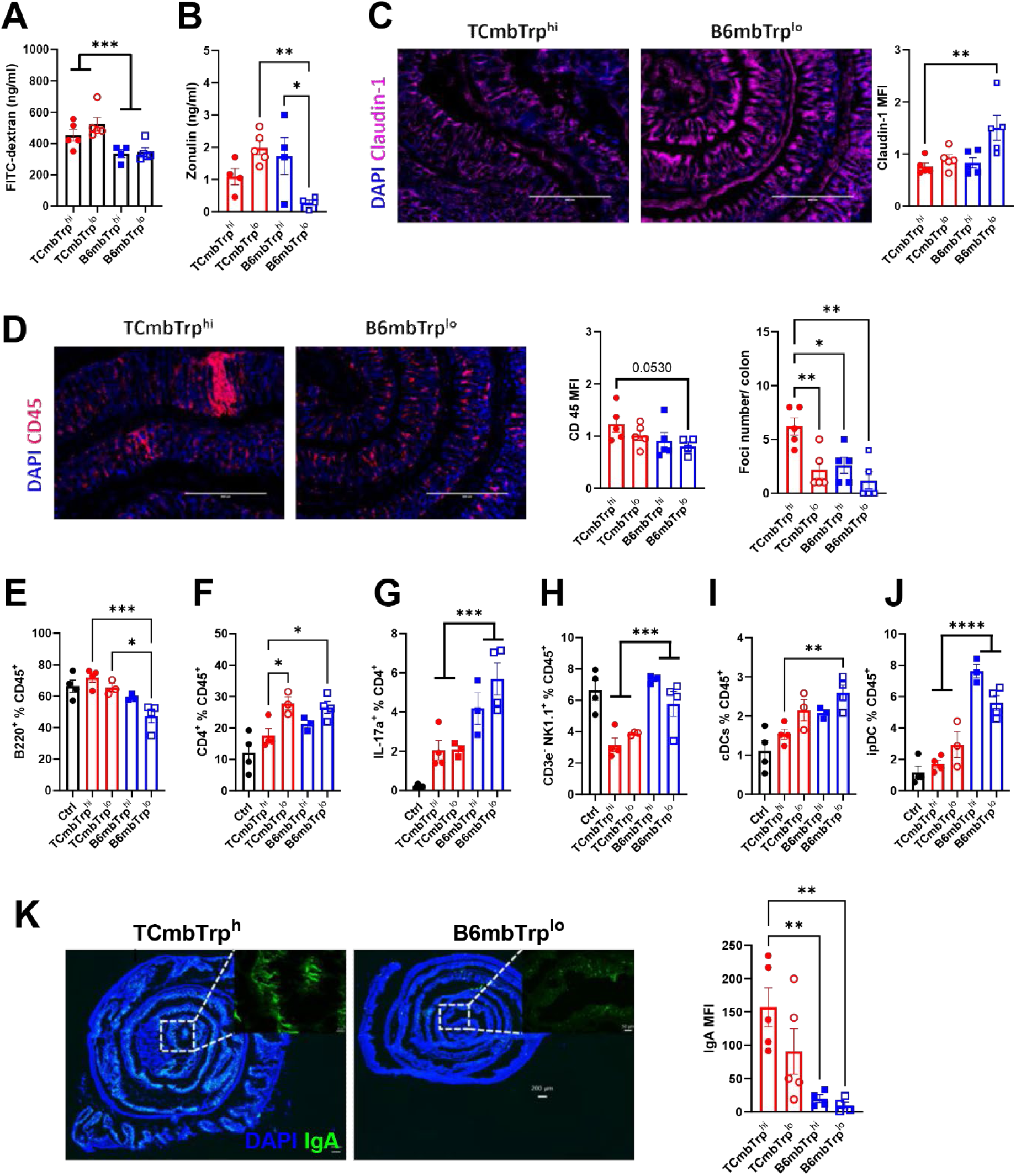
Gut microbiota from lupus-prone mice and high dietary tryptophan disrupt gut barrier integrity and promote intestinal inflammation. (A) Serum FITC-dextran concentration in the four groups of FMT recipient mice after FITC-dextran gavage. (B) Serum zonulin concentrations. (C) Representative images of TCmbTrp^hi^ and B6mbTrp^lo^ ileums stained for Claudin-1 (10X, scale bars: 400 µm) with mean fluorescence intensity (MFI) quantitation shown on the right. (D) Representative images of TCmbTrp^hi^ and B6mbTrp^lo^ colons stained for CD45 (10X, scale bars: 400 µm) with MFI quantitation as well as the number of foci shown on the right. (E – K) Flow cytometry analysis of the colon lamina propia. Frequency of B220^+^ cells (E) and CD4^+^ T cells (F) among CD45^+^ cells. (G) Frequency of IL-17a-producing cells among CD4^+^ T cells. Frequency of CD3e^-^ NK1.1^+^ NK cells (H), cDCs (I) and ipDCS (J) among CD45^+^ cells. (K) Representative images on IgA staining in the colon and IgA signal intensity quantification. N = 4 – 5 per group. Mean + SEM compared with 1-way ANOVA with multiple-comparison tests (B, C, D, E and J), or *t* tests between grouped samples according to the origin of the microbiota (A, G, I and K). *: P < 0.05; **: P < 0.01; ***: P < 0.001; ****: P < 0.0001.

### The combination of lupus gut microbiota and dietary tryptophan resulted in distinct fecal metabolite profiles

Gut microbiota may modulate host immune responses through their metabolites^37^, and we have shown that TC and B6 mice present different fecal metabolite profiles that are altered by dietary tryptophan^14^. Here we used our FMT model to compare the metabolites produced by the TC and B6 gut microbiota developing in the B6 recipients in response to changes in dietary tryptophan. Both positive and negative mode principal component analyses revealed a clear segregation of metabolite profiles between the four groups of FMT recipients (**Figure 3A**). Most of the top 50 metabolites showing a different abundance among the four groups are still unidentified (**Figure S4A-B**). Among the identified features, the tryptophan and several other metabolic pathways were enriched by high tryptophan diet independent of microbiota origins. However, unique pathways distinguished the TC from B6 microbiota recipients fed a high tryptophan diet (**Figure 3B**). Specifically, high tryptophan induced the glycolysis and gluconeogenesis pathway uniquely in TCmbTrp^hi^ mice, while the TCA pathway was enhanced in B6mbTrp^hi^ mice. We have shown that the inhibition of glycolysis with 2-deoxyglucose (2DG) in CD4^+^ T cells and B cells is therapeutic in TC mice^38–40^, and this protective effect can be transferred by FMT from 2DG-treated mice^41^. The current data suggests that high fecal glycolysis may contribute to the enhanced autoimmune activation in TCmbTrp^hi^ hosts. An increased fecal fatty acid oxidation may also contribute to autoimmune activaition in TCmbTrp^hi^ mice, since alterations in fecal lipid metabolism have been reported in SLE patients^42^. Lastly, fecal lipoate metabolism may also contribute to autoimmune activation in TCmbTrp^hi^ mice as lipoic acid has been suggested to have immunomodulatory functions in autoimmune diseases^43^. At the individual metabolite level, high dietary tryptophan also changed the fecal metabolite profiles according the the origin of the microbiota (**Figure 3C**). Among these, TCmbTrp^hi^ mice showed the lowest levels of two short chain fatty acids (SCFA), butyrate and propionate, while high tryptophane had no imoact in B6 FMT recipients (**Figure 3C**; **Figure S4C-D**). SCFAs have well-documented protective effects on gut barrier integrity and autoimmune diseases^44,45^. The abundance of several tryptophan metabolites also segregated with both the microbiota origins and tryptophan diet. Host-produced quinaldic acid, 1-acetylserotonine and 1-methylnicotinamide were higher in TCmbTrp^hi^ mice. In contrast, bacteria-produced 3-dehydroshikimate showed the highest level in B6mbTrp^lo^ mice (**Figure 3C**; **Figure S4E-H**). These metabolites may exert immunomodulatory function on host cells or impact gut microbiome^46^. Overall, factors from both lupus-prone mice microbiota and dietary tryptophan alterated the fecal metabolism including metabolites that may modulate autoimmune activation.

**Figure 3.**
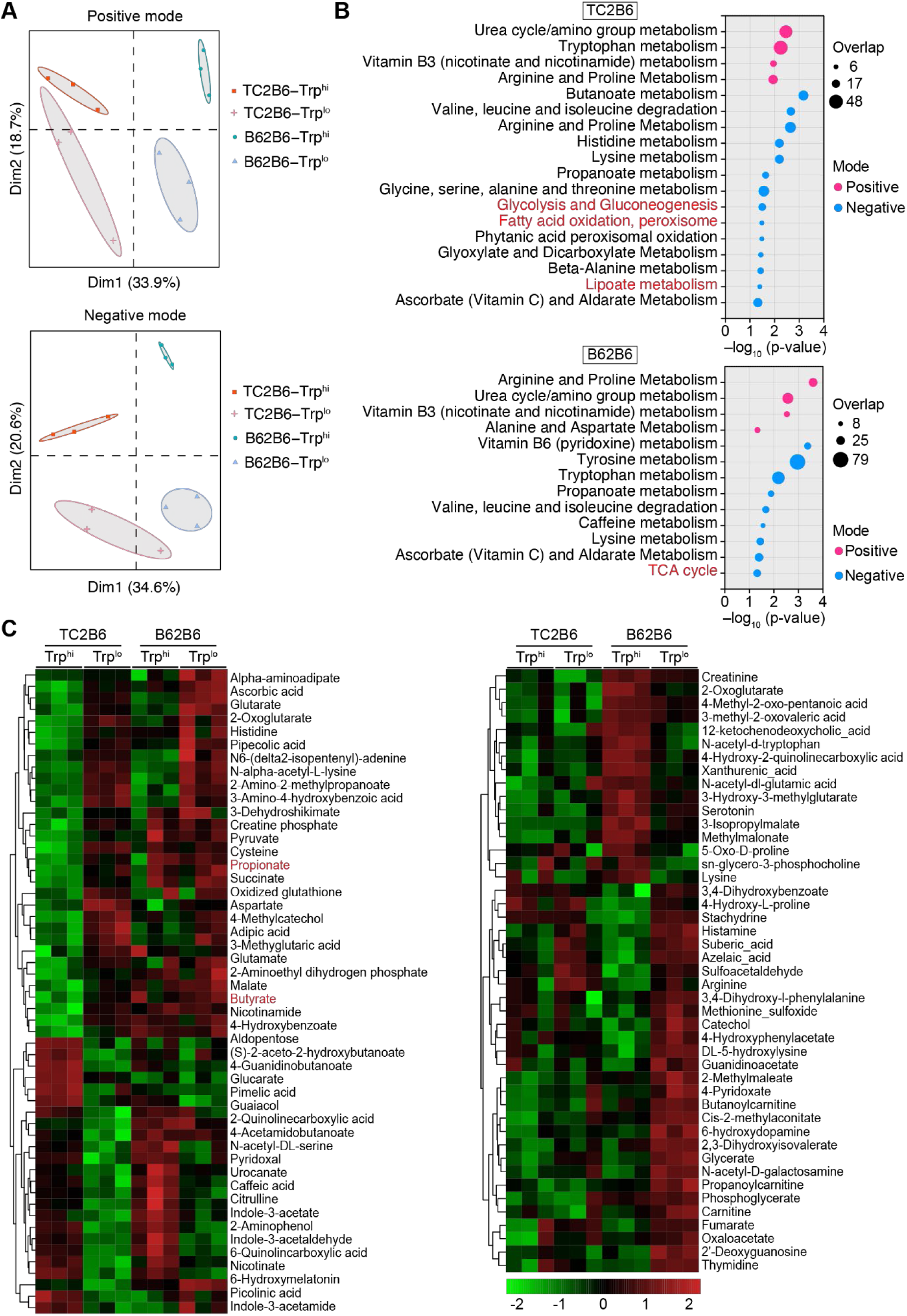
Lupus gut microbiota and dietary tryptophan contribute to distinct fecal metabolite profiles. (A) PCA plots of metabolite features identified by positive and negative ionization. (B) Metabolic pathway analysis of metabolites with significantly differing intensity between tryptophan-high and tryptophan-low groups in B6 mice receiving TC microbiota (TC2B6) or B6 microbiota (B62B6). The pathways unique to each group are labeled with red font (C) Metabolites specifically enriched in TC2B6 or B62B6 mice (overlapping metabolites were not plotted). Left panel: metabolites differentially enriched by tryptophan in TC2B6 mice but not B62B6 mice. Right panel: metabolites differentially enriched by tryptophan in B62B6 mice but not TC2B6 mice. SCFAs are labeled with red font.

### Dietary tryptophan enriched different taxa from the microbiota of TC and B6 mice

To identify bacteria that may contribute to the observed autoimmune activation, we analyzed the fecal microbial composition by 16S rDNA sequencing in a cohort of B6 mice after four weeks of high or low tryptophan diet before and after B6 or TC FMT. To eliminate the contribution of the recipient microbiome, we used germ-free FMT recipients. The post-FMT microbiome diversity was similar between recipient mice that received TC and B6 microbiota, but low tryptophan reduced the diversity in only the recipients of TC FMT (**Figure 4A**). This suggests a different sensitivity to tryptophan between TC and B6 gut microbiota. A principal component analysis showed distinct microbiome profiles for the TCmbTrp^hi^ and B6mbTrp^lo^ groups, with an overlap between the TCmbTrp^lo^ and B6mbTrp^hi^ groups located in between (**Figure 4B**). It should be noted that all recipient profiles were well-separated from their respective donor pools. The relative abundance of 16s rDNA sequencing showed a decreased abundance of *Bacteroidales* in TCmbTrp^lo^ mice while the *Staphylococcaceae* family was uniquely depleted in TCmbTrp^hi^ mice (**Figure 4C**; **Figure S5A-B**). Several species of bacteria were either enriched or depleted by high dietary tryptophan in microbiota of either origin. For instance, high dietary tryptophan enriched the *Bacteroidales* S24-7 and depleted both *Bifidobacterium* and *Clostridium methylpentosum* (**Figure 4D**; **Figure S5C-E)** in both TC and B6 microbiota. However, dietary tryptophan also enriched specific taxa, such as *Coriobacteriaceae, Lachnospiraceae*, *Bacteroides*, *Rikenellaceae*, *Parabacteroides distasonis* and *Streptococcaceae bacterium* RF32, depending on the origin of the microbiota (**Figure 4D**; **Figure S5F-K**). Noticeably, the TCmbTrp^hi^ samples contained the highest number of taxa with a significantly different abundance, 7 out of 10 of which were uniquely amplified by the combination of TC origin and high tryptophan*. Ruminoccocus gnavus* (*Rg)* featured preeminently among these taxa (**Figure 4E**), which was confirmed by qPCR analysis in an independent cohort of FMT recipients (**Figure 4F**). The qPCR analysis also showed the dual effect of the microbiota origin and dietary tryptophan, with *Rg* abundance being higher in TCmbTrp^lo^ than in B6mbTrp^hi^ mice, and a similar trend between B6mbTrp^hi^ and B6mbTrp^lo^ mice. 16S rDNA sequence analysis cannot distinguish the LP-producing *Rg* pathobiont strain, *Rg*2, from non-LP producing strain *Rg1*^47^. However, the growth of *Rg2,* but not *Rg1,* was enhanced by supplemental tryptophan *in vitro* (**Figure 4G-H**), suggesting that dietary tryptophan expand the *Rg2* population present in the TC microbiota. Overall, these results suggest that both dietary tryptophan and the origin of the microbiota shape the architecture of gut microbiome. This applies specifically to *Rg*, a group of bacteria now firmly associated with severe lupus ^10^, which was found at a greater abundance in recipient microbiota of TC origin and was further enriched by high tryptophan.

**Figure 4.**
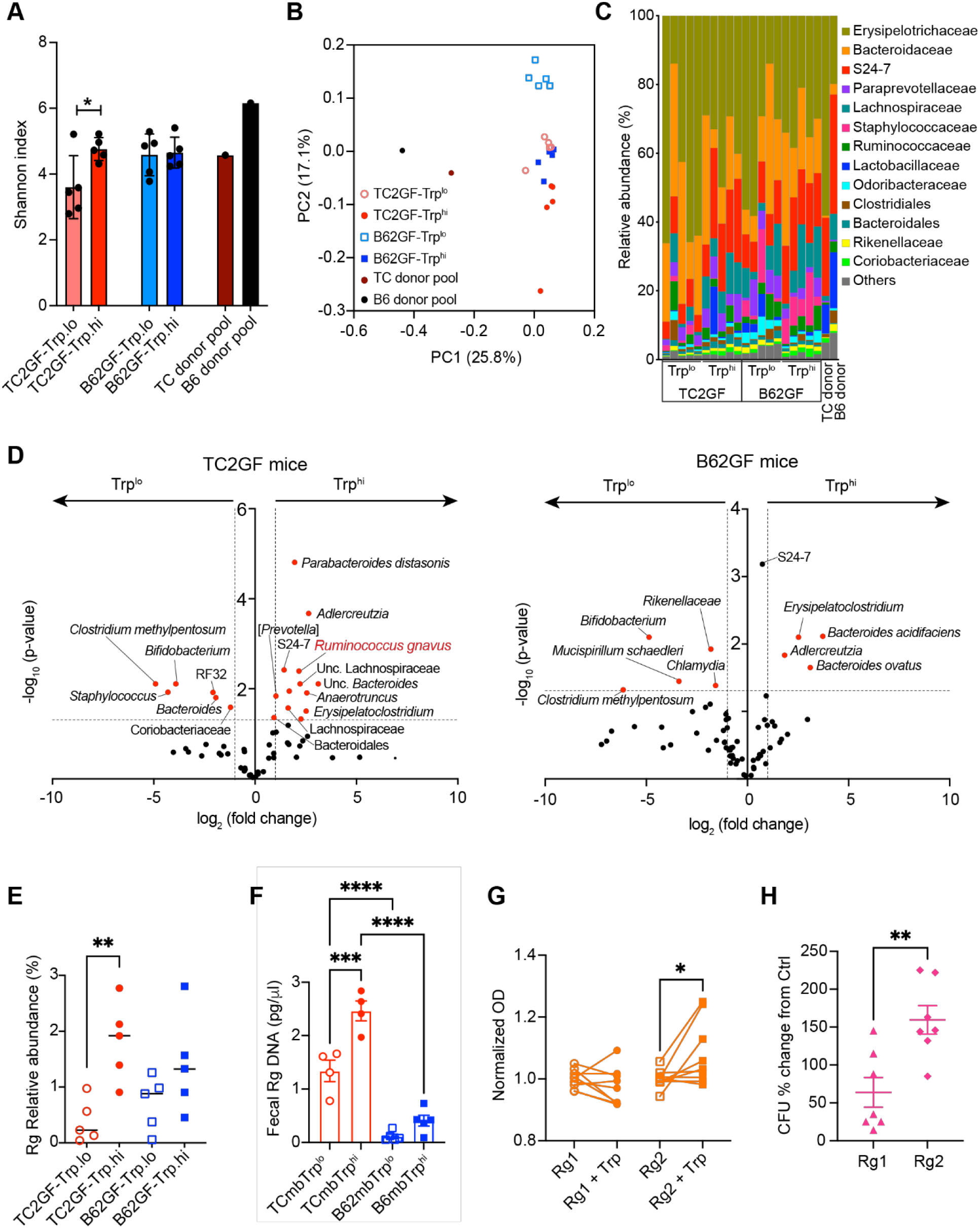
Dietary tryptophan enriched different taxa in the microbiota from TC and B6 mice. (A) Gut microbiome diversity in recipient and pooled donor groups. (B) Unweighted Unifrac PcoA plots for 16sDNA sequencing analysis. (C) 16S DNA sequencing family abundance. (D) Volcano plots showing the taxa differentially enriched by tryptophan in B6 mice receiving TC microbiota (TC2B6, top) or B6 microbiota (B62B6, bottom). *Ruminoccocus gnavus* (Rg) relative abundance in the four groups of recipients measured by 16S DNA sequencing (E) and qPCR (F). (G) OD600 of *Rg*1 or *Rg*2 cultured *in vitro* with or without 100 μM L-tryptophan 4 h post inoculation. Results from 3 cohorts with 3 – 4 replicates each were normalized to the mean values without tryptophan for each cohort as 1. (H) Colony forming unit (CFU) changes induced by 2 mM L-tryptophan on *Rg*1 or *Rg*2 *in-vitro* cultures at 6 h post inoculation. Mean + SEM compared with t tests (A and H), 1-way ANOVA with multiple-comparison tests (E and F), or paired t test (G). **: P < 0.01; ***: P < 0.001; ****: P < 0.0001.

### Fecal *Rg* is enriched in lupus prone TC mice

The greater abundance of *Rg* in the recipients of TC microbiota suggested that these bacteria are enriched in TC mice. Since *Rg* blooms correspond to disease flares in SLE patients ^10^, we speculated that *Rg* populations may expand with disease progression in TC mice. To test this hypothesis, we longitudinally tested *Rg* abundance in the feces of age-matched TC and B6 mice housed side by side, in two cohorts each housed in separate rooms. In both cohorts, TC and B6 mice had similar *Rg* levels in the first 3 months of age (**Figure 5A**), when TC mice do not produce yet autoantibodies. After 3 months, the *Rg* population expanded in TC mice, while remaining relatively constant in B6 mice (**Figure 5A**). One cohort also presented a late *Rg* bloom in B6 mice and a contraction in TC mice. Interestingly, a lower *Rg* level was detected in TC.Rag1^-/-^ mice, which lack mature T and B cells (**Figure 5B**), suggesting that adaptive immunity is necessary for the expansion of *Rg* in lupus-prone mice. Next, we assessed whether *Rg* abundance correlated with host autoimmune phenotypes in a cohort of 6 - 9 months old TC mice in which *Rg* abundance varied (**Table S1**). The levels of serum anti-dsDNA IgA and anti-dsDNA IgM were significantly higher in the mice with a fecal *Rg* abundance above the median level, and a similar trend was observed for anti-dsDNA IgG (**Figure 5C-E**). Furthermore, serum IL-6 and IFN-γ levels were positively correlated with fecal *Rg* abundance in these mice (**Figure 5F-G**). Overall, these results suggest that *Rg* expands in TC mice as they start producing autoantibodies, and *Rg* abundance is positively correlated with autoimmune activation markers.

**Figure 5.**
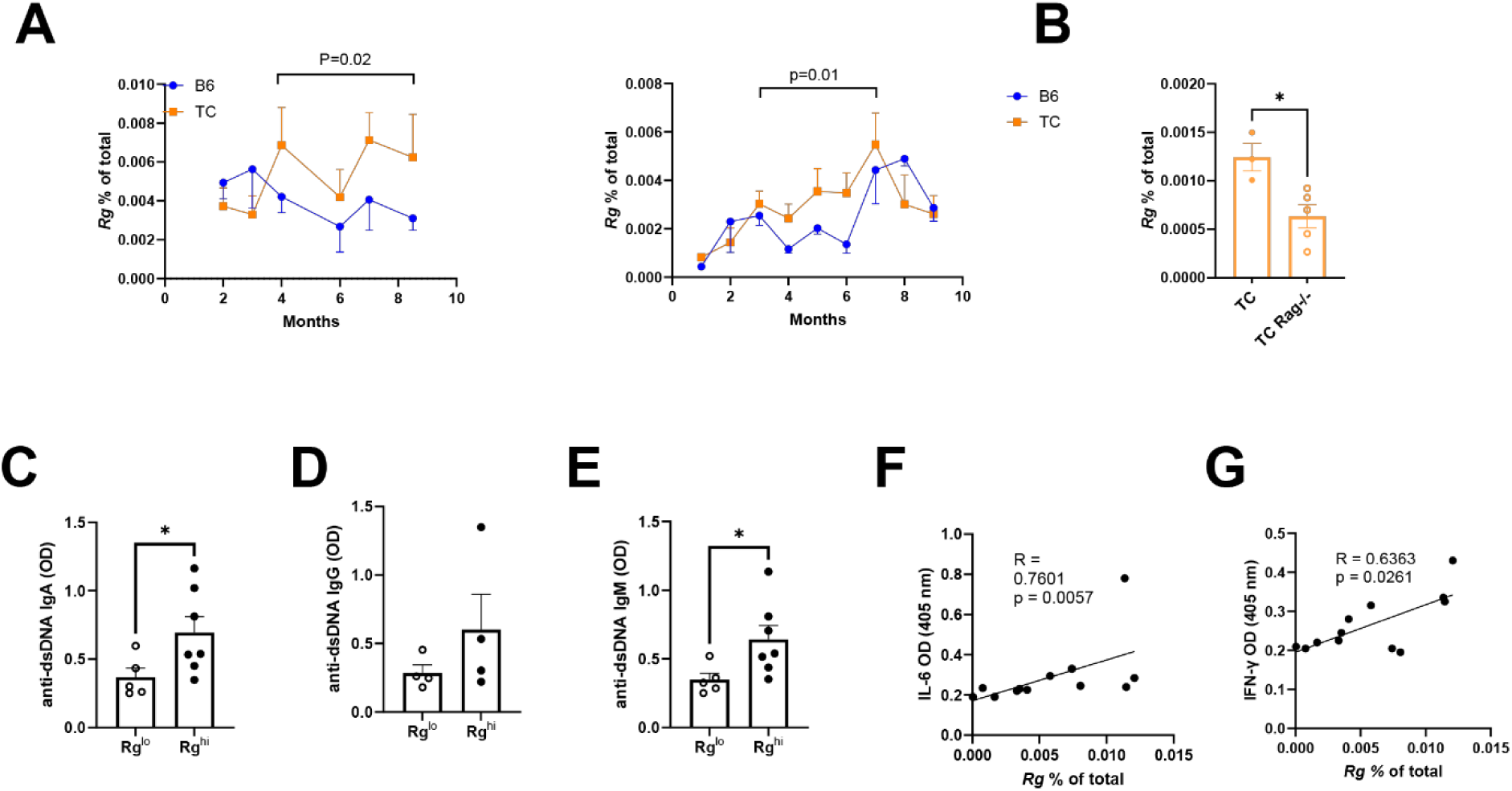
*Rg* is enriched in lupus prone mice. (A) Longitudinal quantification of fecal *Rg* abundance in two cohorts of TC and B6 mice (N = 4 - 5 mice per strain in each). The brackets correspond to the time periods in which the two strains were significantly different by 2-Way ANOVA. (B) Fecal *Rg* levels in 3-month-old TC and TC.Rag1^-/-^ mice compared by t test. (C – G) A cohort of 6 – 9-month-old TC mice was assessed for fecal *Rg* levels and serum anti-dsDNA IgA (C), anti-dsDNA IgG (D) and anti-dsDNA IgM (E), IL-6 (F) and IFN-γ (G). For anti-dsDNA antibodies (C – E), mice were distributed in Rg^lo^ and Rg^hi^ relative to the median level and compared with Mann-Whitney tests. IL-6 correlation was evaluated with the Spearman test and IFN-γ with a Pearson test. Mean + SEM. *: P < 0.05.

### Fecal *Rg* promoted the development of lupus in TC mice

To test the role of *Rg* in the development of lupus in TC mice, we continuously treated mice with metronidazole starting at 2 months of age (**Figure 6A**). Fecal *Rg* was depleted after 1 month treatment, without affecting the global bacterial content (**Figure 6B-C**). After 5 - 7 months of treatment, *Rg* depletion from gut microbiota was associated with a reduction of anti-dsDNA IgG production, splenomegaly, CD4^+^ T cell expansion, as well as a decreased ratio of follicular helper T (Tfh) to follicular regulatory T (Tfr) cells with a relative expansion of Tfr cells (**Figure 6D-G; Figure S6A-B**), all of which are markers of disease activity in lupus. On the other hand, *Rg* depletion increased CD25 and FOXP3 expression by Treg cells (**Figure 6H-J**). Treg expressing low levels of CD25 and FOXP3 have a reduced suppressive function, and are expanded in SLE^48^ ^49^. The frequency of Treg cells was also increased in the mLN of treated mice (**Figure 6K**), suggesting a direct expansion at sites closer to the microbiota. These results suggest that depletion of *Rg* from TC gut microbiota suppress autoimmune activation and expands functional Treg cell populations.

**Figure 6.**
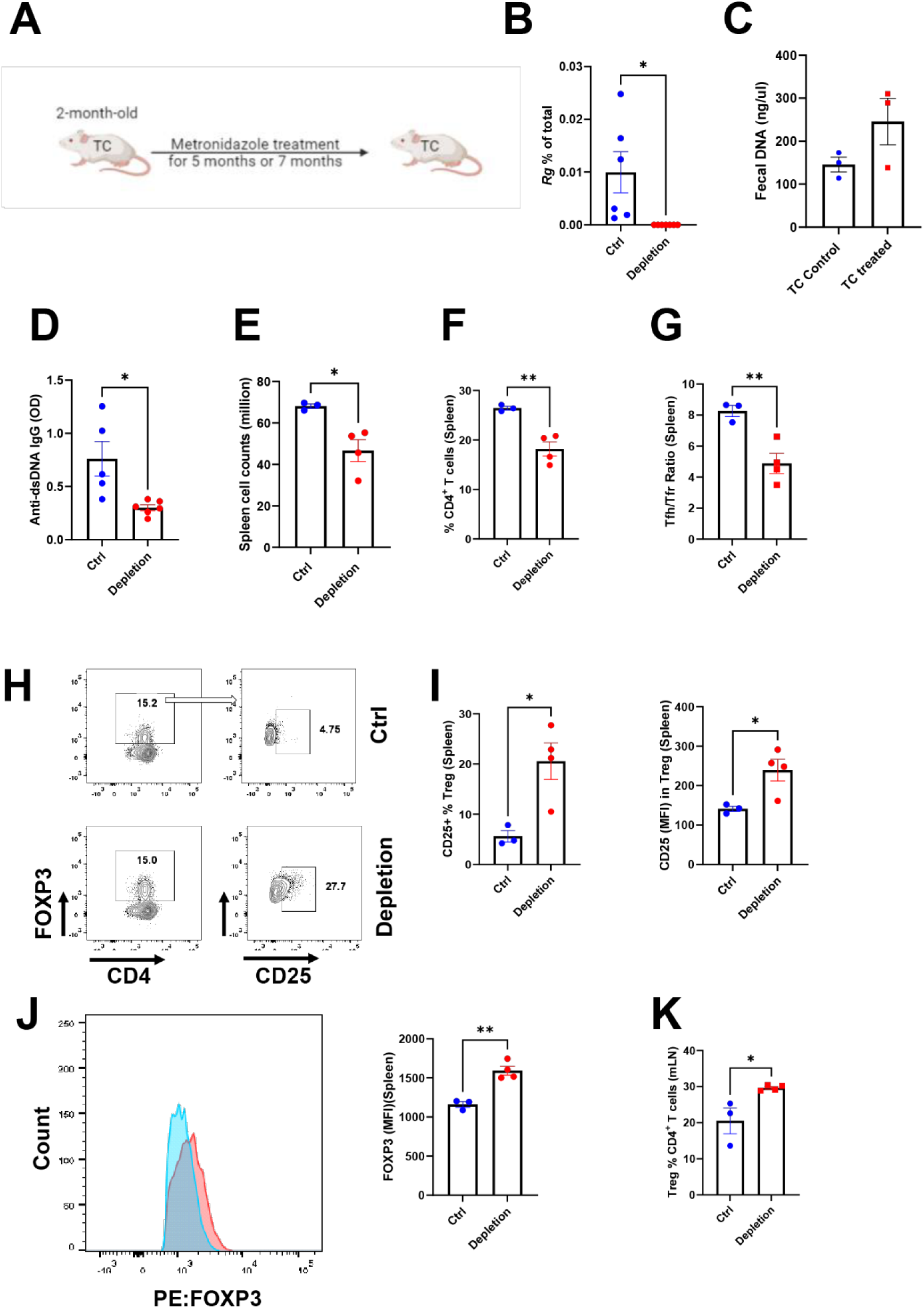
*Rg* depletion reduced autoantibody production and expanded Treg cells. (A) Experimental design for *Rg* depletion in TC mice by metronidazole. *Rg* levels (B) and total fecal microbial DNA level (C) in control and metronidazole-treated TC mice. (D) Serum anti-dsDNA IgG. (E) Splenocyte counts. (F) Frequency of splenic CD4^+^ T cells. (G) Splenic Tfh/Tfr ratio. (H - J) Frequency of splenic CD25^high^ Treg cells, CD25 and FOXP3 expression by Treg cells, with representative FACS plots showing CD25 (H) and FOXP3 (J) expression. (K) Treg cell frequency out of CD4^+^ T cells in mLN. Mean + SEM compared with t tests. *: P < 0.05; **: P < 0.01.

To further evaluate the role of *Rg* in autoimmune activation, we colonized antibiotic-treated B6 and TC mice with the LP-producing *Rg* pathobiont strain, *Rg*2^47^ or the control *Rg1* (**Figure 7A**). *Rg* was detected in fecal samples one month after colonization of TC mice with *Rg2* (**Figure 7B**), while *Rg*1 showed a significant lower level of colonization (**Figure S7A).** Attempts to colonize three independent cohorts of B6 mice with *Rg2* were unsuccessful (**Figure S7B**). *Rg*2 colonization of TC mice increased the production of anti-dsDNA IgG (**Figure 7C**) and the frequency of activated CD69^+^ CD4^+^ T cells and effector memory CD4^+^ T cells, while it decreased the frequency of naïve CD4^+^ T cells (**Figure 7D-F**). The frequency of IFN-γ producing CD4^+^ T cells and IL-10 producing non-regulatory T cells, two cell types expanded in SLE progression^50^ ^51^, were also increased in *Rg*2 monocolonized mice (**Figure 7G-H**). Additionally, the frequency of germinal center B cells was increased (**Figure 7I**). *Rg2*-colonized TC mice also produced anti-lipoglycan IgG which was positively correlated with the level of serum anti-dsDNA IgG (**Figure 7J-K**), confirming the association of *Rg* cell wall LP with anti-dsDNA IgG production that was reported in SLE patients^47^. Taken together, our results suggest that *Rg2* expansion which occurs preferentially in TC mice, induced autoimmune activation.

**Figure 7.**
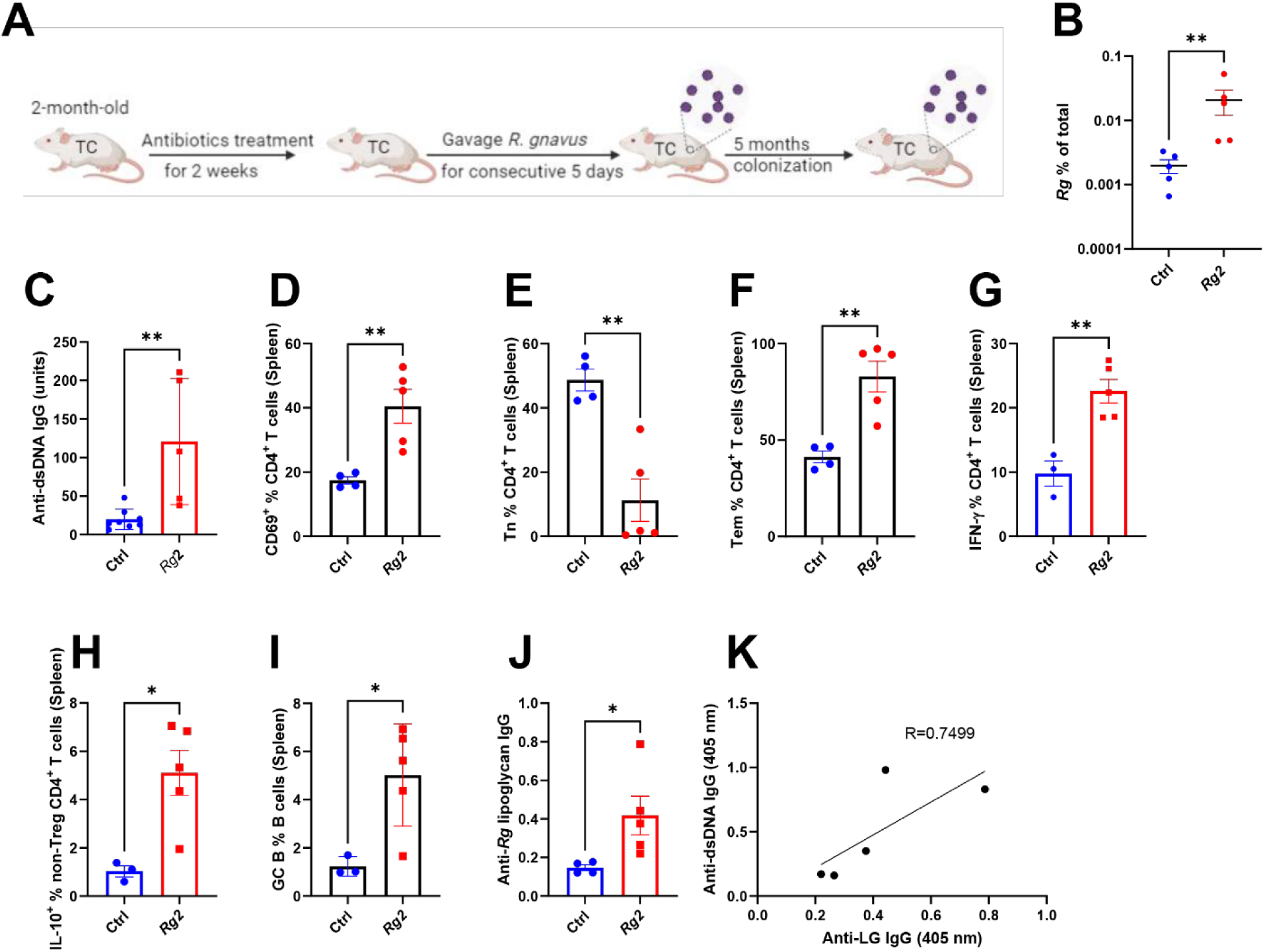
*Rg2* monocolonization induced autoimmune activation in TC mice. (A) Experimental design. (B) *Rg* levels in control and *Rg* mono-colonized TC mice post 1 month inoculation. Immune phenotypes were analyzed 5 months after colonization. (C) Serum anti-dsDNA IgG. Frequency of splenic CD69^+^ CD4^+^ T cells (D), naïve CD4^+^ T cells (E), effector memory CD4^+^ T cells (F), IFN-γ^+^ CD4^+^ T cells (G), IL-10-producing non-Treg T cells (H), and germinal center B cells (I). (J) Serum anti-*Rg* lipoglycan IgG. (K) Correlation between serum anti-lipoglycan IgG and anti-dsDNA IgG. Mean + SEM compared with t tests. *: P < 0.05; **: P < 0.01; ***: P < 0.001; ****: P < 0.0001.

### *Rg2* induced Treg cell apoptosis

*Rg2* colonization increased effector T cell functions, and *Rg* depletion enhanced FOXP3 expression on Treg cells in TC mice. We validated the effect of *Rg2* on T cells by showing an increased frequency of proliferating Ki67^+^ FOXP3^neg^non-Treg CD4^+^ T cells that was not significant for Treg cells (**Figure 8A**), and a decreased frequency of FOXP3^+^ CD4^+^ Treg cells (**Figure 8B**) in non-autoimmune B6 mLN cells exposed to *Rg* lysates. Further, a higher level of activated caspase 3 was detected in these *Rg*2 stimulated Treg cells, which was not detected in non-Treg cells (**Figure 8C**), suggesting a pro-apoptosis function from *Rg*2 on Treg cells. However, LP-negative *Rg1* showed less pronounced effects. Taken together, *Rg*2 reduced the frequency of Treg cells and increased conventional helper T cell proliferation, possibly as a consequence of reduced Treg suppression.

**Figure 8.**
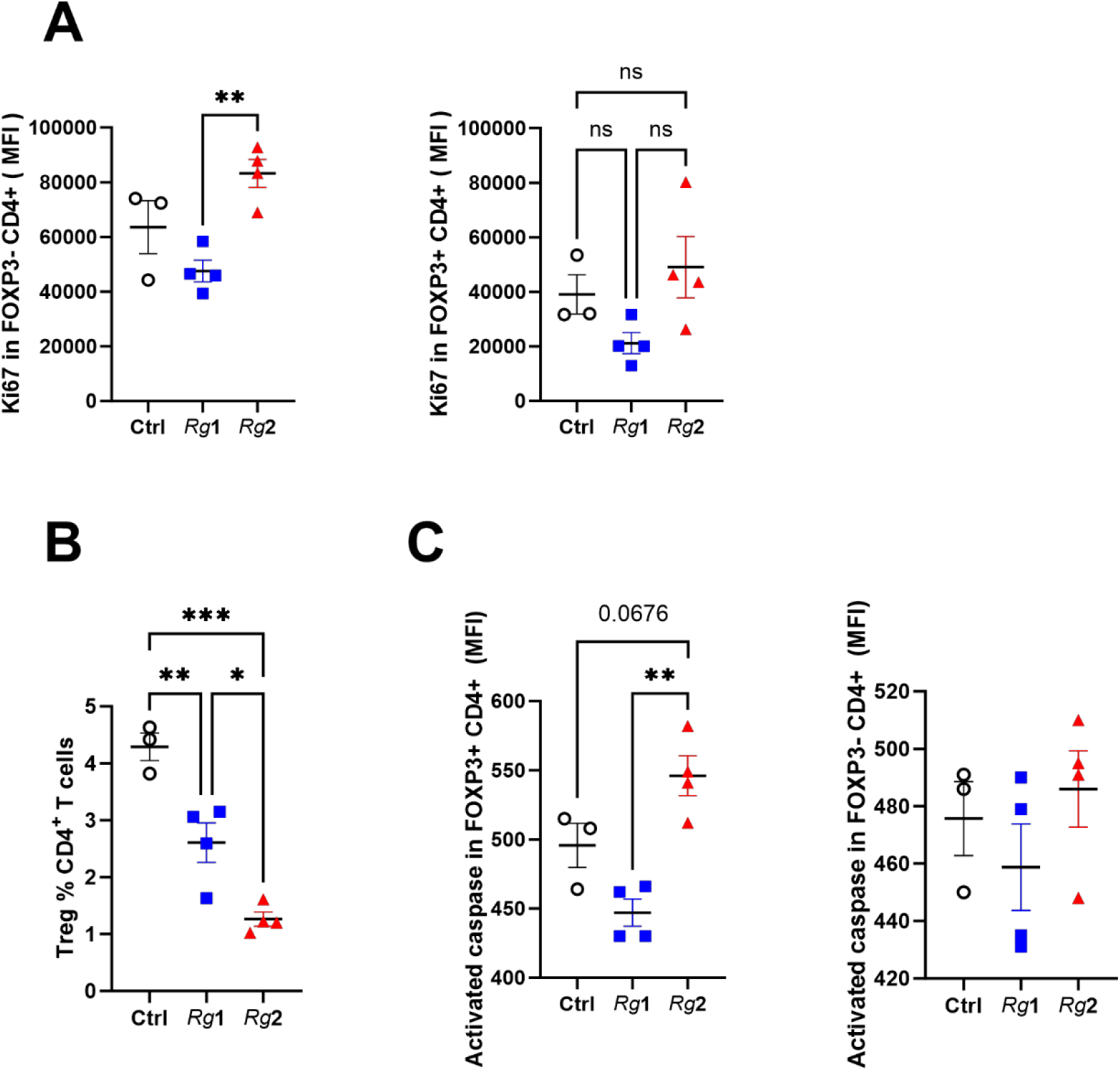
*Rg* modulates CD4^+^ T cell populations. **B6** mLN cells were exposed to *Rg*1 or *Rg2* bacterial lysate for 72 h. (A) Ki67 expression in non-Treg CD4^+^ T cells and in Treg cells. (B) Treg cell frequency in CD4^+^ T cells. (C) Activated Caspase 3 levels in Treg and non-Treg T cells. Mean + SEM compared with t tests. *: P < 0.05; **: P < 0.01; ***: P < 0.001.

## Discussion

Gut microbiome alterations, which have been reported in lupus patients and mouse models ^52,53^, may modify the host immune system through direct interactions between immune cells and bacteria or bacterial metabolites that impact immune cell metabolism and functions^54^. We have shown that restricting dietary tryptophan changed the composition of the gut microbiome in the TC lupus-prone model and improved disease outcome^14^. Further, we have shown that variations in dietary tryptophan changed not only the phenotypes of CD4^+^ T cells from TC mice^14^, but also their metabolism, at least partially in an intrinsic manner, in a way that was different from changes in the T cells from B6 controls^15^. This suggested that genetic factors controlled the skewed tryptophan metabolism in TC mice as genetic susceptibility controls their immune system. To further analyze the complex interactions between genetic susceptibility, the immune system, the microbiome and tryptophan metabolism, we used a reductionist experimental system in which the only variables were the origin of the microbiota and the amount of dietary tryptophan. We transferred the gut flora from TC mice or B6 mice into B6 mice fed with high or low tryptophan chow, therefore eliminating genetically driven autoimmune activation. The combination of FMT with microbiota of TC origin and high dietary tryptophan increased autoimmune responses. The production of autoantibodies, the expansion of splenocytes, the frequency of B cells, as well as the infiltration of the kidneys with T cells were enhanced in TCmbTrp^hi^ mice. Conversely, these mice showed a reduced frequency of mLN Treg cells. These results suggest that the combinations of factors within the gut microbiota of lupus prone mice and high dietary tryptophan promoted autoimmune activation of a non-autoimmune immune system. The gut tissue where intestinal microbiota encounters dietary tryptophan displayed an inflammation with a differential distribution of immune cells as well as a decreased barrier integrity in TCmbTrp^hi^ mice. Dietary tryptophan altered the fecal metabolomic profiles as well as the bacterial distribution depending on the FMT origin. Low dietary tryptophan reduced the diversity of the microbiota of TC but not B6 origin, suggesting a higher frequency of tryptophan-sensitive bacteria in the TC gut microbiota. Dietary tryptophan also expanded or reduced specific bacterial taxa in the gut microbiome of TC and B6 origins. *Rg*, a bacterium expanded during flares in SLE patients^9^, was uniquely enriched in TCmbTrp^hi^ mice. *Rg* is also pro-inflammatory in Crohn’s disease through its glucorhamman polysaccharide ^55^.

To investigate the possible pathogenic role of *Rg* in lupus development, we first compared its abundance in TC and B6 mice in a longitudinal study. *Rg* spontaneously expanded in the gut of TC mice after 3 months of age, when autoimmune activation accelerates while it remained low in B6 mice. The depletion of *Rg* from TC mice with oral metronidazole improved disease outcomes. Specifically, *Rg* depletion alleviated splenomegaly and autoantibody production, decreased the frequency of CD4^+^ T cells and plasma cells but increased the frequency of Treg cells. *Rg* is sensitive to metronidazole ^56,57^; but this antibiotic also globally depletes anaerobes. To specifically test the pathogenic role of *Rg*, we monoclonized antibiotic-treated TC mice with *Rg2,* the LP-producing strain that is close to the LP-producing *Rg* strains that are expanded in human SLE ^10,47^. Consistent with the FMT results, *Rg2* stable monocolonization was successful in TC but not in B6 mice. Healthy GF B6 mice were successfully colonized with *Rg2* by vertical transmission^47^, The difference in outcomes may be qualitative due to the continuous exposure of neonates to their parents’ *Rg2* for 21 days instead of the one week gavages performed in this study, or due to a greater ability for *Rg2* to colonize the gut from neonates relative to adult mice. Nonetheless, these results suggest that *Rg2* expands preferentially in genetically driven immune inflammation in TC mice, corresponding to the blooms observed in SLE patients with high disease activity ^9,10^.

We were also not able to achieve a stable colonization of TC mice with the non-LP producing *Rg1*. *Rg1* cannot be distinguished from *Rg2* with either 16S rDNA PCR or sequencing. Therefore, we cannot assess the relative contribution of Rg2 to the *Rg* expansion found in the TC microbiota, either at steady state or after FMT with high dietary tryptophan. The respective requirements of *Rg1* and *Rg2* to establish and maintain a stable population are at the present unclear, but the enhanced growth of *Rg2* but not *Rg1* in response to high tryptophan may contribute to its ability to colonize TC mice. I also suggest that *Rg2* may preferentially expand *in vivo* in TCmbTrp^hi^ mice and contribute to autoimmune activation and their impaired intestinal barrier.

We showed that *Rg* depletion in TC mice was associated with an increased frequency of FOXP3^+^ CD4^+^ Treg cells with a higher CD25 and FOXP3 expression, suggesting enhanced suppressive function. A specific effect of *Rg2* on Treg cells was established by showing a higher apoptosis of non-autoimmune Treg cells exposed to *Rg2* lysate. Strikingly, non-autoimmune mLN helper CD4^+^ T also showed a higher proliferation when stimulated with *Rg*2, which could be a direct effect or an indirect effect through Treg depletion. *Rg2* colonization of TC mice enhanced their levels of serum anti-dsDNA IgG in correlation with anti-*Rg* lipoglycan IgG, as well as T cell activation, with increased frequencies of effector T cells, IFN-γ producing and IL-10 producing non-Treg CD4^+^ cells, as well as GC B cells. These results suggest a direct role of *Rg2* in autoimmune activation in TC mice. Overall, this study showed a synergistic pathogenic effect between the gut microbiome that develops in an autoimmune environment and dietary tryptophan. Gut dysbiosis develops with autoimmunity in TC mice since young pre-autoimmune mice show a similar microbiota than congenic B6 controls that was not able to induce autoimmune activation through FMT^14^. Here we showed that once established, lupus gut dysbiosis activates the immune system and impairs intestinal integrity in healthy mice with high tryptophan availability. This synergistic effect includes the expansion of *Rg2*, a human lupus pathobiont, which enhances autoimmune activation in lupus-prone mice. *Rg* population expands in TC mice at the time they start developing autoimmunity, but cannot stably colonize B6 mice, suggesting that an activated immune system is necessary to create a permissive environment, which is further enhanced by dietary tryptophan.

Tryptophan has now been implicated in exacerbating three autoimmune diseases through the gut microbiome in mouse models, lupus^14,15^, multiple sclerosis^58,59^ and rheumatoid arthritis (RA)^60^. Tryptophan metabolites have been shown to possess immunomodulatory functions^46^. For example, kynurenine activates the aryl hydrocarbon receptor (AhR), a transcription factor that controls local and systemic immune responses, in dendritic cells, leading to TGF-β production^61^. In addition, tryptamine ameliorated symptoms of experimental autoimmune encephalomyelitis (EAE) by activating AhR and suppressing neuroinflammation^62^. On the other hand, microbiota-produced kynurenic acid expanded Th17-inducing GPR35^+^Ly6C^+^ macrophages^59^ and indole promoted the differentiation of Th17 cells that exacerbates collagen-induced arthritis^60^. In lupus-prone mice, tryptamine, which is more abundant than in congenic controls, works as a pro-inflammatory factor by promoting mTOR pathway activation and glycolysis in CD4^+^ T cells^15^. Interestingly, we found in the present study an enhanced glycolytic pathway in the fecal metabolites of TCmbTrp^hi^ mice. We also found several kynurenine pathway metabolites that were enriched in the gut of in TCmbTrp^hi^ mice, including quinolinic acid, quinaldic acid, 1-methylnicotinamide and 6-hydroxynicotinate. Elevated levels of quinaldic acid in the synovial fluid of RA and osteoarthritis (OA) patients correlated with disease activity^63^. The increased level of quinaldic acid in the TCmbTrp^hi^ gut may also have pro-inflammatory effects on immune cells and/or intestinal epithelial cells, skewing gut homeostasis. Other tryptophan metabolites may also contribute to inflammation, individually or in combination. High dietary tryptophan also reduced the production of butyrate and propionate by the microbiota of TC origin, which may contribute to an inflamed gut environment in TCmbTrp^hi^ mice, considering the beneficial effects of SCFAs in gut homeostasis^44,45^. In addition, SCFAs promote intestinal Treg differentiation^64,65^, and SCFAs induced by dietary fiber were associated with a reduced gut dysbiosis and disease activity in a TLR7-induced model of lupus ^7^. These results suggest that tryptophan was differently metabolized by TC or B6 mice microbiota and the availability of tryptophan also modified other metabolic pathways. The differential distribution of tryptophan metabolites may work synergistically with other metabolites to create a pro-inflammatory gut environment.

Taken together, our study indicates that high levels of dietary tryptophan expand specific bacterial taxa and their metabolites from TC gut microbiota that induce autoimmune activation and intestinal inflammation independently from the immune system. These bacterial taxa include *Rg2*, a proposed pathobiont presented by SLE patients with high disease activity. Depletion and monocolonization experiments showed that the lipoglycan-producing *Rg2* strain accelerated autoantibody production and T cell activation. The expanded *Rg2* may increase intestinal inflammation and enhance systemic immune responses via dampened gut integrity and bacterial translocation. Our results also suggest a direct effect of Rg2 on CD4^+^ T cells.

There are some limitations in this study. We have not isolated *Rg* from fecal samples of TC mice and therefore identified the specific *Rg* strains that are present either at steady state or expanded by tryptophan in the FMT experiment. *Rg* present in TC mice may be *Rg*2 or a mixture of several different strains. In the FMT assay, other bacteria besides *Rg* also showed correlations to microbiota origins or tryptophan diets. We have not tested the possible pathogenic impact of each additional species on autoimmune activation. Fecal metabolites that showed different levels between microbiota origins or tryptophan diets may also contribute to phenotypes we observed in recipients. These metabolites should be evaluated *in-vitro* and/or *in-vivo* on lupus pathogenesis. We demonstrated that *Rg*2 affects helper T cell proliferation as well as Treg development and function. However, if this reaction is *Rg*2 antigen-specific, like lipoglycan, is unknown.

## Material and Methods

### Study design

The study aimed to directly test the hypothesis that changes in dietary tryptophan alter the host gut microbiome from lupus-prone mice, which indirectly modulates autoimmune activation. For this purpose, we eliminated the genetic factors underlying autoimmune activation and potential variations in intrinsic tryptophan metabolism, and we compared the effect of high and low dietary tryptophan on the gut microbiota from TC or B6 mice in the same non-autoreactive immune system. To accomplish this, we performed FMT from lupus-prone TC and congenic healthy control B6 mice into antibiotic-treated or gnotobiotic B6 recipients, which were fed with high or low tryptophan. Autoimmune phenotypes were evaluated in the FMT recipients, including serum and fecal autoantibodies, immunophenotyping by flow cytometry of the spleen and intestines, and histology. Fecal metabolites were compared between B6 and TC FMT recipients with an untargeted screen using LC-HRMS/MS. The distribution of fecal bacteria was compared between FMT recipient mice by 16S rDNA sequencing. We then focused on the role of *Ruminococcus gnavus* (*Rg*), a bacteria taxa that was selectively amplified by high tryptophan in the recipients of TC microbiome. The abundance of fecal *Rg* was compared in two longitudinal cohorts of B6 and TC mice. Immune phenotypes were evaluated in TC mice that were treated with metronidazole, which depletes *Rg*, and controls, and in TC mice monocolonized with *Rg*. Finally, the effect of *Rg* on immune cells was evaluated *in vitro*.

### Mice and treatment

C57Bl/6J (B6) mice were originally purchased from the Jackson laboratory. B6.*Sle1.Sle2.Sle3* (TC) and TC.*Rag*1^-/-^ mice have been described previously^66,67^. All mice were bred and maintained at either the University of Florida (UF) or University of Texas Health Science Center at San Antonio (UTHSCSA) in specific pathogen-free (SPF) conditions. Gnotobiotic (GF) B6 mice were obtained from the UF GF facility and used as FMT recipients in SPF conditions as previously described^14^. Only female mice were used with age-matched controls for each experiment. Mice were strictly housed with littermates or with mice from the same strain within the same age group (i.e. less than 4 weeks apart). This housing policy was to avoid transfers of autoimmune activation or attenuation that occur between B6 and TC mice that share their microbiome^14^. Unless specified, mice were fed with irradiated Envigo 7912 standard chow. 7 to 10-month-old TC mice with high serum anti-dsDNA IgG titers and age-matched B6 mice were used as FMT donors. 15 fecal pellets were pooled from each group of donor mice and homogenized in 3.75 mL sterile PBS containing 0.5g/L L-cysteine. Homogenates were centrifuged at 1500 rpm for 8 minutes, and 200 uL of supernatant were gavaged into recipient mice, which were 8-week-old B6 mice that were either GF or pre-treated with a cocktail of antibiotics [AMNV: ampicillin (0.05%; Fisher Scientific AC61177-0050), metronidazole (0.05%; Fisher Scientific AC210340050), neomycin (0.05%; Cayman 14287), and vancomycin (0.025 %; Cayman 15327) in the drinking water for 2 weeks before transfer. The recipients were fed with high- or low-tryptophan chows for 4 weeks, with 1 week before FMT and 3 weeks after FMT. Briefly, the chemically defined chows (Research Diets) differ only by their tryptophan content, which is 1.19% for high tryptophan. Low tryptophan was achieved by a tryptophan-free chow on weekdays supplemented by 0.19% tryptophan on week-end days, corresponding to an approximate overall 0.03% tryptophan^14^. Each cohort contained 4 groups of mice of recipient mice. Before euthanasia, gut permeability was evaluated by gavaging mice that were fasted for 4 h with 5 mg FITC-dextran 4000 (Sigma -Aldrich) in 200 μl PBS. FITC-dextran was quantified in the serum 1.5 h later by detecting fluorescence using the SpectraMax M5/M5e Multimode Plate Reader. To deplete *Rg*, 2-month-old TC mice were continuously treated with 0.15 % metronidazole in drinking water for 5 - 7 months, and *Rg* depletion was tested in fecal samples by qPCR after 1 month treatment. For *Rg2* (CC55_001C NIH) and *Rg1* (29149 ATCC) monocolonization, 2-month-old TC and B6 mice were treated with AMNV for 2 weeks. Two days after the termination of AMNV treatment, mice were colonized by daily oral gavage with10^8^ CFU of *Rg2* in 200 μL of sterile PBS for 5 consecutive days. The study was carried out in accordance with the guidelines from the Guide for the Care and Use of Laboratory Animals of the Animal Welfare Act and the National Institutes of Health. All animal protocols were approved by the Institutional Animal Care and Use Committee at UF and UTHSCSA.

### Bacterial assays and culture, *in-vitro* growth and *ex-vivo* exposure to immune cells

Fecal pellets were collected and stored at −80°C until DNA extraction with DNeasy PowerLyzer PowerSoil Kit (Qiagen; 12855-100). Fecal *Rg* abundance was determined with *Rg*-specific 16S rDNA primers: Forward: 5’-GGACTGCATTTGGAACTGTCAG-3’ Reverse: 5’-AACGTCAGTCATCGTCCAGAAAG-3’ (ref. ^47^). Serial dilutions of *Rg2* DNA were used to generate a qPCR standard curve. A total of 100 ng fecal DNA was used for each test qPCR reaction, from which the abundance of *Rg* was presented as percentage of total DNA. Before and after AMNV treatment, the amount of total fecal bacteria was measured by qPCR with 16S rDNA specific primers: Forward: 5’-ACTCCTACGGGAGGCAGCAGT -3’, and Reverse: 5’-ATTACCGCGGCTGCTGGC -3’ (ref. ^68^). Based on the formula “*Copy number per μl = (ng/μl × [6.022×10*^23^*]) / (length × [1×10*^9^*] ×650)*” and the length of *Rg* genome (∼3550000 bp), the *Rg*2 qPCR standard curve was used to calculate copy numbers from Ct values.

*Rg*1 (29149 ATCC) and *Rg*2 (CC55_001C NIH) were streaked on blood agar (TSA with 5% sheep blood) plates, then individual colonies were grown in 5 ml of BHI media (Anaerobe Systems) under anaerobic conditions for 16 h. Cultures were diluted down to 0.02 OD600. To test the effect of tryptophan on Rg growth, 100 μM or 2mM tryptophan was added into culture, and growth was evaluated either as OD600 of the cultures at 4 h, or CFU at 6 h. To prepare bacterial lysates, *Rg*1 and *Rg*2 were cultured for 6 h to collect bacteria at log phase. The OD600 was measured to calculate the CFUs based on growth curves of *Rg*1 and *Rg*2. The bacterial pellets were washed twice with PBS and then reconstituted in 1 mL PBS. Bacteria were sonicated with Branson Digital Sonifier at 60% 3 times for 30 seconds. For the bacterial lysate stimulation assay, 2 x 10^5^ mLN cells from B6 mice were cultured in 100 uL complete RPMI medium (Corning, 10-040-CV) with 1 μg/mL anti-CD3 (BD Bioscience; BDB553057) and 1 μg/mL anti-CD28 antibodies (BD Biosciences BDB553294). Bacterial lysates were added into the culture with a 1:10 ratio of mLN cells to bacteria. The phenotypes of CD4^+^ T cells were evaluated by flow cytometry after 72 h exposure.

### Renal and intestinal tissue histology

Kidneys were embedded in OCT medium (Fisher Scientific) and snap-frozen at − 80 °C. Ileum and colon were prepared as “Swiss rolls” before being embedded in OCT medium and snap-frozen. 7 µm thick sections were mounted on histology slides set at room temperature for 20 min before being placed into PBS to dissolve the OCT. The sections were then fixed with cold acetone for 10 min, washed 3 times, and blocked with 10% normal rat serum (Equitech-Bio) in PBS for 30 min. Renal tissue sections were stained with anti-CD3e-APC (1:50; eBioscience 145-2C11). The ileum sections were stained with rabbit anti-Claudin-1 (1:100 dilution; Invitrogen MH25) followed by goat anti-rabbit IgG-AF700 (1:2000; Invitrogen A-21038). The colon sections were stained with anti-CD45-APC/Cyanine7 (1:25 dilution; Biolegend 30-F11) and anti-IgA-FITC (1:50 dilution; BD Biosciences). Fluorescence intensity was analyzed using ImageJ.

### Antibody and cytokines measurements

Serum and fecal anti-dsDNA IgG, IgM and IgA were quantified by ELISA with 1:100 dilutions as previously described^14^. Anti-dsDNA IgG were also visualized on *Crithidia luciliae* slides (Bio-Rad) with serum diluted 1:10 followed by an incubation with goat anti-mouse IgG-FITC (1:100; Southern Biotech). Images were acquired on an inverted fluorescence microscope (Olympus). The corrected total cell fluorescence (https://theolb.readthedocs.io/en/latest/#) from each *Crithidia luciliae* cell was quantified with ImageJ. IL-6 or IFN-γ were quantitated with BD Biosciences ELISA kits in serum samples diluted 1:50 assayed in duplicate. The absorbance was detected with the Promega GloMax® Explorer microplate reader at 450 nm.

### Flow cytometry

Single-cell suspensions were isolated from the gut using the lamina propria dissociation kit with gentleMACS tissue dissociator (Miltenyi Biotech) or from spleen and mLN by smashing tissues thoroughly between frosted glass slides. Cells were stained in 2.5% FBS and 0.05% sodium azide in PBS. Fluorochrome-conjugated antibodies used in this study are listed in Supplementary Table 2. Dead cells were excluded with fixable viability dyes (eFluor780 or LIVE/DEAD™ Fixable Yellow Dead Cell Stain Kit; Thermo Fisher Scientific). Intracellular staining was performed with a fixation/permeabilization kit (eBioscience). For cytokine detection, splenocytes were stimulated with the Leukocyte Activation Cocktail (BD Biosciences) at 37 ^0^C for 4 h. All samples were acquired on an LSR Fortessa flow cytometer (BD Biosciences) and analyzed with FlowJo software. The gating strategies are shown in Figure S2-S3.

### 16sRNA gene sequencing

Fecal samples stored at −80°C were processed for microbiota sequence analysis. 16S rRNA libraires were constructed and sequenced as described previously ^68^. Briefly, a minimal sequence depth of 44,966 2 x 300-bp reads (paired-end) per sample was obtained and processed using QIIME 2 (v2020.8). Reads were merged, quality trimmed, and clustered into operational taxonomic units at 97% sequence similarity. Taxonomy was assigned using Greengenes 13_8.

### Fecal metabolomics

An untargeted analysis of the fecal metabolome was conducted as previously described ^15^. Briefly, samples homogenized in 5 mM of ammonium formate were normalized to the lowest protein concentration, and spiked with internal standards (IS) consisting of L-tryptophan-2,3,3-D_3_ (40 μg/mL) and 4 g/mL of creatine-D_3_, leucine-D_10_, L-tyrosine Ring-^13^C^6^, L-leucine-^13^C^6^, L-phenylalanine^13^C_6_, N-BOC-L-tert-leucine, N-BOC-L-aspartic acid, succinic acid-2,2,3,3-D_4_, salicylic acid D_6_, and caffeine-D_3_. Metabolites were extracted by protein precipitation with a solution of 8/1/1 (v/v/v) Acetonitrile/ Methanol/ Acetone. Samples were centrifuged at 20,000 x g for 10 min at 4°C to pellet the protein. Supernatants transferred into Eppendorf tubes were dried under a gentle stream of nitrogen at 30°C. The dried extracts were re-suspended with 25 μL reconstitution solution consisting of 10 ug/mL injection standards (BOC-L-Tyrosine, BOC-L-Tryptophan and BOC-D-Phenylalanine). Resuspension was allowed at 4°C for 10-15 min then samples were centrifuged at 20,000 x g for 10 min at 4°C. Supernatants were collected into LC-vials for LC-MS analysis. Global metabolomics were performed using high-resolution mass spectrometry coupled with ultra-high performance liquid chromatography (UHPLC) analyzed in positive and negative heated electrospray ionization as previously described ^14^. Separation was achieved on an ACE 18-PFP 100 x 2.1 mm, 2 μm column using a gradient with mobile phase A as 0.1% formic acid in water and mobile phase B as acetonitrile with a column temperature of 25°C. Gradient elution was ramped from 0% B to 80% B over 13 min at 350 μL/min, which increased to 600 μL/min between 16.80 and 17.50 min for column flush and re-equilibration. The runtime was 20.50 min and full scan at 35,000 mass resolution was acquired from 2 μL injection in positive and 4 μL injection in negative ion mode. After normalizing to total ion chromatogram, intensities were tested for group significance using unpaired Student *t*-test. Metabolites from this method were identified as they are matched with retention time and *m/z* value to a reference standard.

### Statistics

Statistical analyses were performed with the Graphpad Prism 9.0 software. Differences between groups were evaluated by one-way ANOVA with correction for multiple tests, unpaired t tests or Mann-Whitney nonparametric test, as indicated in the text. Unless specified, all tests are two-tailed. To test correlations between two variables, Pearson or Spearman tests were used. Results were expressed as means ± standard error of the mean (s.e.m.). The levels of statistical significance were set at *: *P* < 0.05, **: *P* < 0.01, ***: *P* < 0.001 and ****: *P* < 0.0001.

## Acknowledgments

We thank Dr. Tim Garrett and the staff of the UF SECIM from the metabolomic analysis, as well as the staff from the Flow Cytometry Shared Resource at UTHSA.

