## Supplemental tables and figures for "Dietary tryptophan and genetic susceptibility expand gut microbiota that promote systemic autoimmune activation"

**Table S1. Partition of 6 - 9 months old TC mice according to their fecal *Rg* levels.**

| ***Rg***$^{low}$ | ***Rg***$^{high}$ |
|---|---|
| 0.1208035 | 0.0579602 |
| 0.1147157 | 0.0408034 |
| 0.1136032 | 0.0352198 |
| 0.0805393 | 0.0332125 |
| 0.0764159 | 0.0165211 |
| 0.0741981 | 0.0003436 |

Mice were partitioned into $Rg^{low}$ and $Rg^{high}$ according to their fecal *Rg* levels shown for each mouse in ng/ml, relative to the group median.

**Supplementary Table 2. Antibodies information**

| Target Antigen | Conjugation | Clone | vendor | Catalog # | Dilution |
|---|---|---|---|---|---|
| CD3ε | BV711 | 17A2 | BioLegend | 100241 | 1:100 |
| BCL-6 | Alexa Fluor 647 | K112-91 | BD Biosciences | 561525 | 1:100 |
| CD4 | FITC | RM4-5 | BD Biosciences | 553047 | 1:100 |
| CD25 (IL-2Rα) | AF700 | PC61.5 | eBioscience | 56-0251 | 1:100 |
| CD44 | PE | IM7 | BioLegend | 103008 | 1:100 |
| CD45.2 | BV650 | 104 | Biolegend | 109835 | 1:100 |
| CD45R (B220) | PE-CF594 | RA3-6B2 | BD Biosciences | 562290 | 1:100 |
| CD62L | APC | MEL-14 | BD Biosciences | 553152 | 1:100 |
| CD69 | PE-Cy7 | H1.2F3 | BioLegend | 104512 | 1:100 |
| CD95 | Biotin | Jo2 | BD Biosciences | 554256 | 1:100 |
| CD11b | Per-CP | M1/70 | BD Biosciences | 550993 | 1:100 |
| CD11c | PE-Cy7 | HL3 | BD Biosciences | 558079 | 1:100 |
| CD19 | PB | eBio1D3 | eBioscience | 48-0193-82 | 1:100 |
| CD8a | AF700 | 53-6.7 | BD Biosciences | 100729 | 1:100 |
| Ly6G | PE | 1A8 | BD Biosciences | 551461 | 1:100 |
| CD138 | APC | 281-2 | BioLegend | 142506 | 1:100 |
| CD185 (CXCR5) | Purified | 2G8 | BD Biosciences | 560577 | 1:100 |
| CD279 (PD-1) | eFluor405 | RMP1-30 | Thermo Fisher | 13-9981 | 1:100 |
| FOXP3 | FITC | FJK-16S | Thermo Fisher | 11-5773 | 1:100 |
| GL7 | eFluor450 | GL7 | Thermo Fisher | 48-5902 | 1:100 |
| IFN-γ | Pacific Blue | XMG1.2 | BioLegend | 505818 | 1:100 |
| IL-10 | PE-Cy7 | JES5-16E3 | BioLegend | 505026 | 1:100 |
| Ki-67 | PE-Cy7 | SolA15 | Thermo Fisher | 25-5698 | 1:100 |
| Caspase 3 | BV450 | C92-605 | BD Biosciences | 560627 | 1:100 |
| Siglec-F | BV421 | E50-2440 | BD Biosciences | 562681 | 1:100 |
| MHC II | PE | M5/114.15.2 | BioLegend | 107608 | 1:100 |
| NK1.1 | PE-Cy7 | PK136 | BioLegend | 108714 | 1:100 |
| CD49b | PerCP/Cy5.5 | Hma2 | BioLegend | 103520 | 1:100 |
| IL-17 | PE | TC11-18h10.1 | BioLegend | 506904 | 1:100 |
| F4/80 | APC | BM8 | BioLegend | 123116 | 1:100 |
| PDCA-1 | PE | eBio927 | eBioscience | 12-3172-81 | 1:100 |

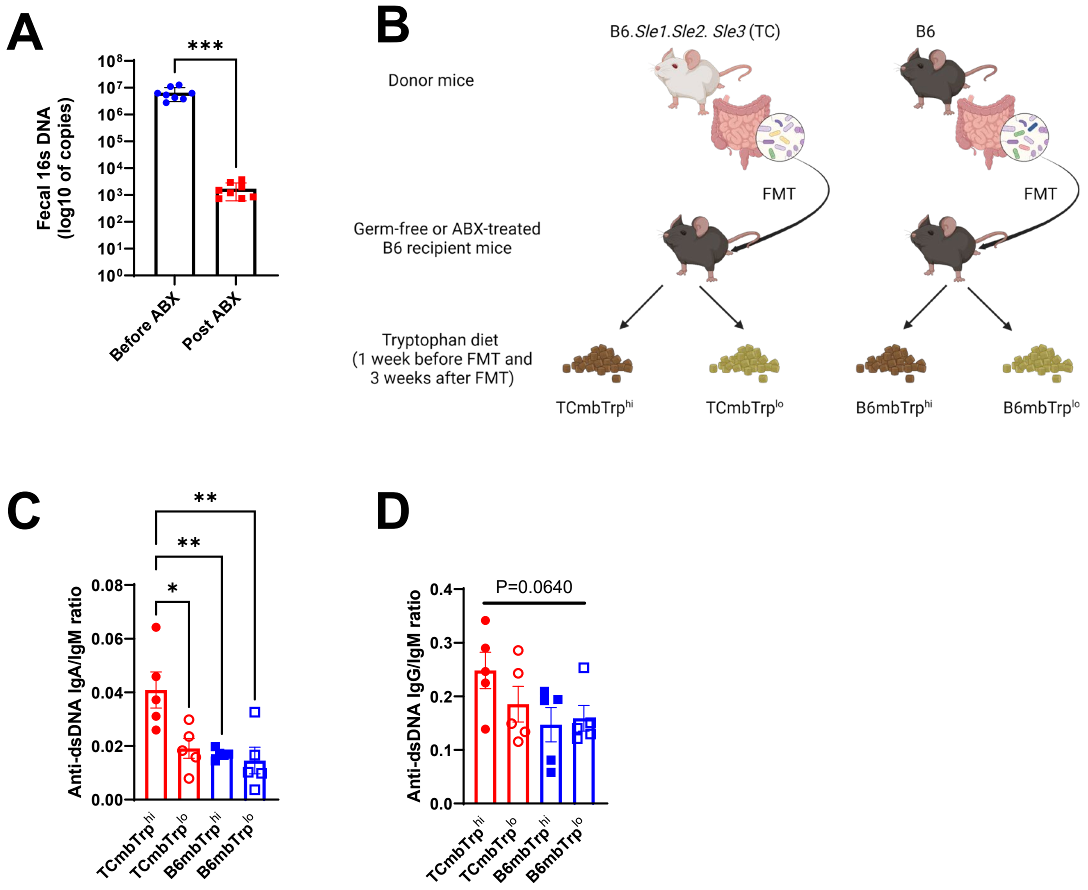


**Figure S1. Both lupus-prone mice gut microbiota and high dietary tryptophan promote autoimmune activation.** (A) 16s DNA copy number in the gut microbiome of B6 mice before or after broad-spectrum antibiotic (ABX) treatment. (B) FMT experimental design. Serum anti-dsDNA IgA / IgM (C) and anti-dsDNA IgG / IgM (D) ratio. Mean + SEM compared with a paired t test (A) and with 1-way ANOVA with multiple-comparison tests (C and D). *: P < 0.05; **: P < 0.01.

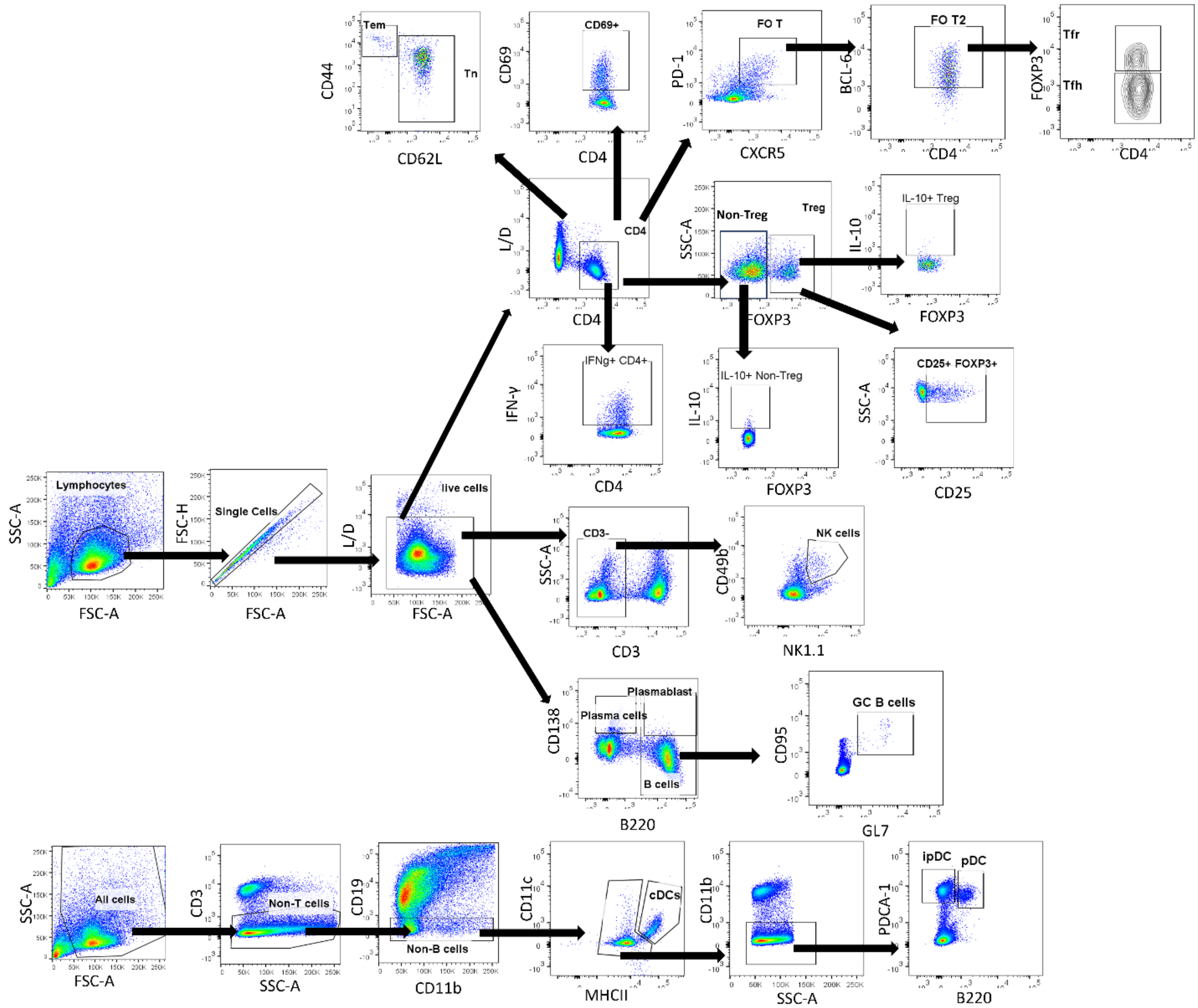


**Figure S2. Representative FACS gating strategy for different cell populations in splenocytes.**

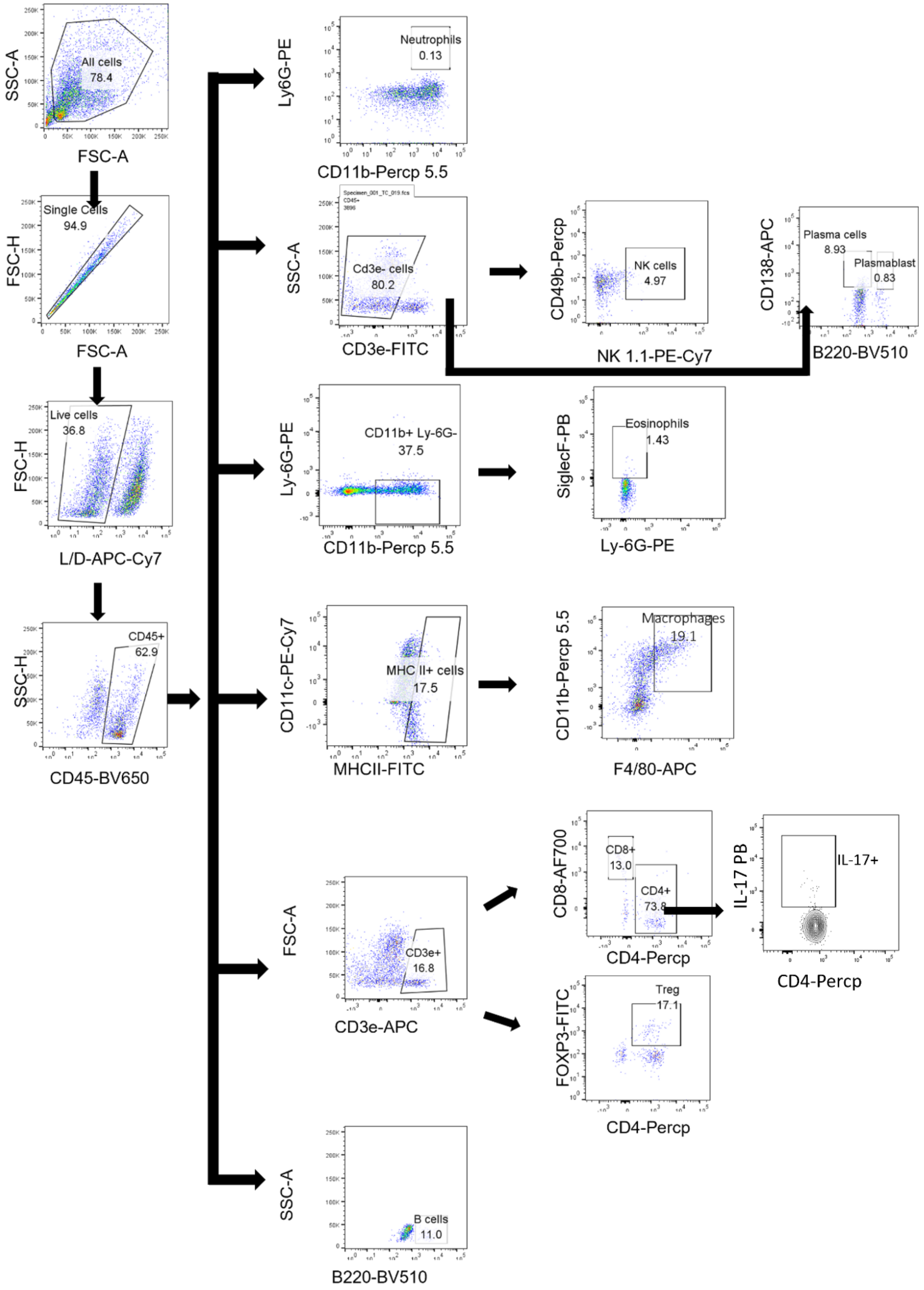


**Figure S3. Representative FACS gating strategy for different cell populations in colon lamina propria cells.**

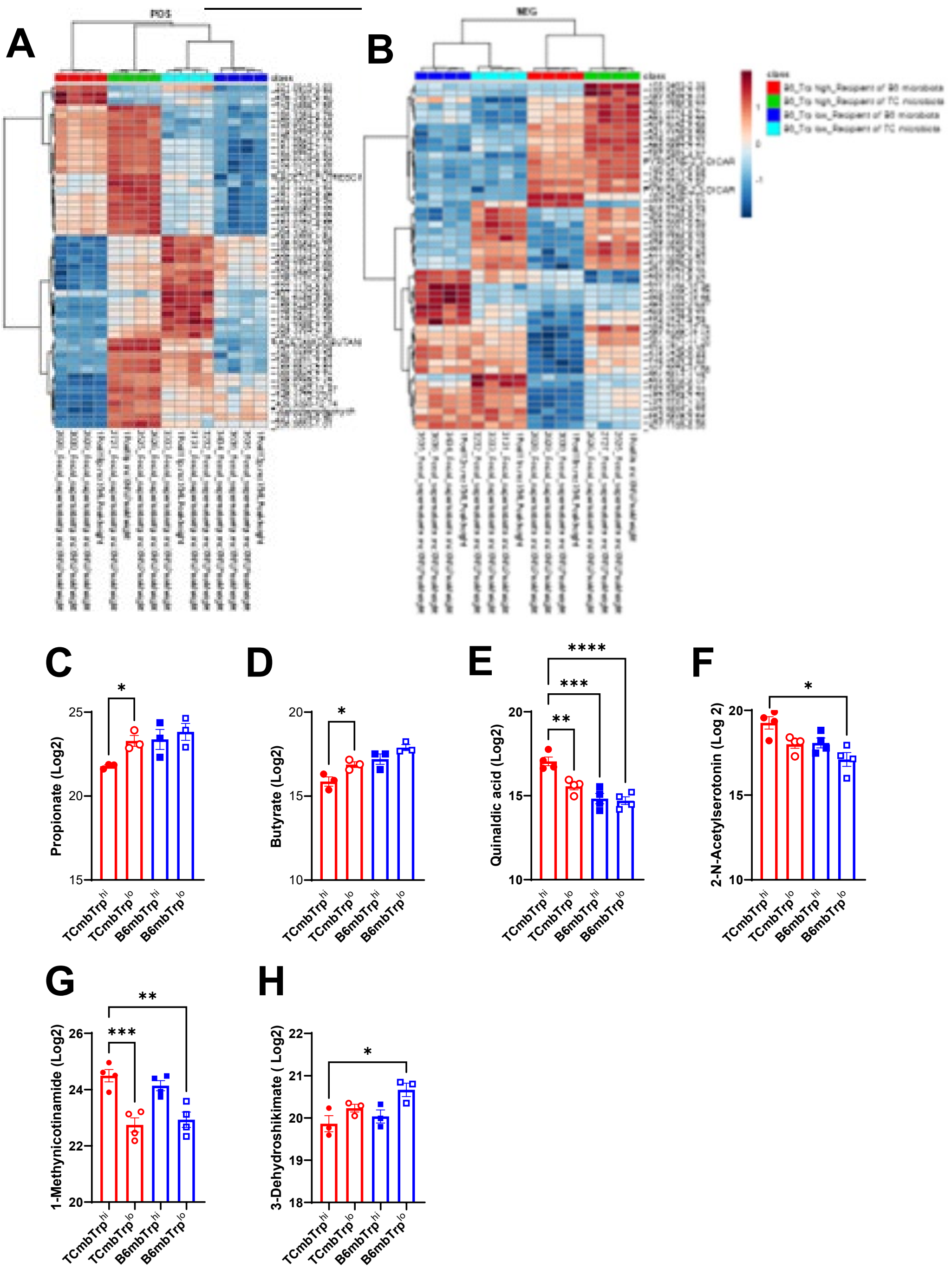


**Figure S4. Lupus gut microbiota and dietary tryptophan contribute to distinct fecal metabolite profiles.** Top 50 metabolites showing a statistically significant different abundance between 4 groups under positive (A) and negative (B) modes. Abundance of propionate (C), (D) butyrate, quinaldic acid (E), 1-acetylserotonine (F), 1-methylnicotinamide (G) and 3-dehydroshikimate (H) in the 4 groups. Mean + SEM compared with 1-way ANOVA with multiple-comparison tests. *: $P < 0.05$; **: $P < 0.01$; ***: $P < 0.001$; ****: $P < 0.0001$.

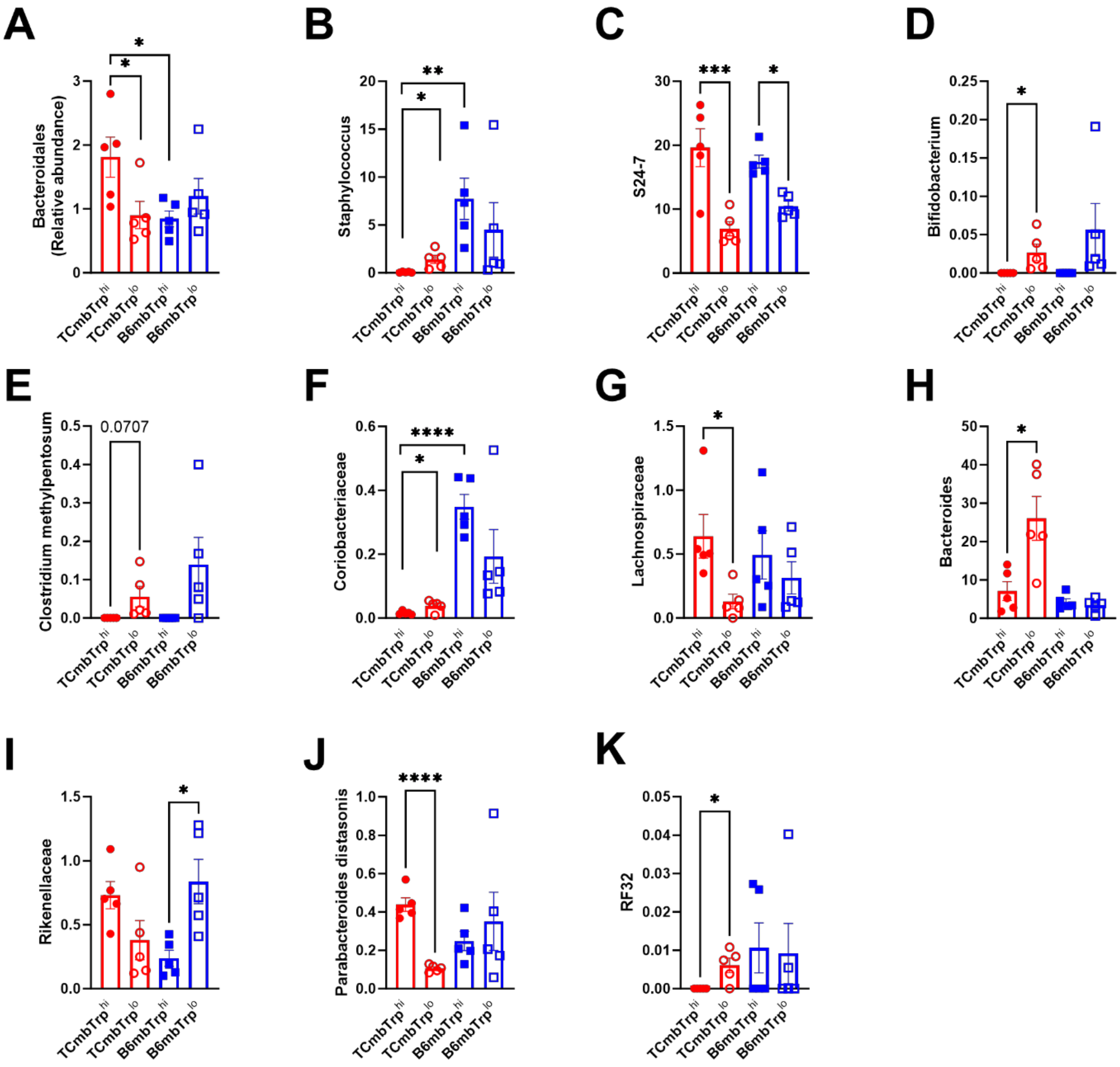


**Figure S5. Relative abundance of bacteria taxa affected by microbiome and/or dietary tryptophan.** *Bacteroidales* (A), *Staphylococcaceae* (B), *Bacteroidales* S24-7 (C), *Bifidobacterium* (D); *Clostridium methylpentosum* (E), *Coriobacteriaceae* (F), *Lachnospiraceae* (G), *Bacteroides (H)*, *Rikenellaceae* (I), *Parabacteroides distasonis* (J) and *Streptococcaceae* bacterium RF32 (K). Mean + SEM compared with 1-way ANOVA with multiple-comparison tests. *: $P < 0.05$; **: $P < 0.01$; ***: $P < 0.001$; ****: $P < 0.0001$.

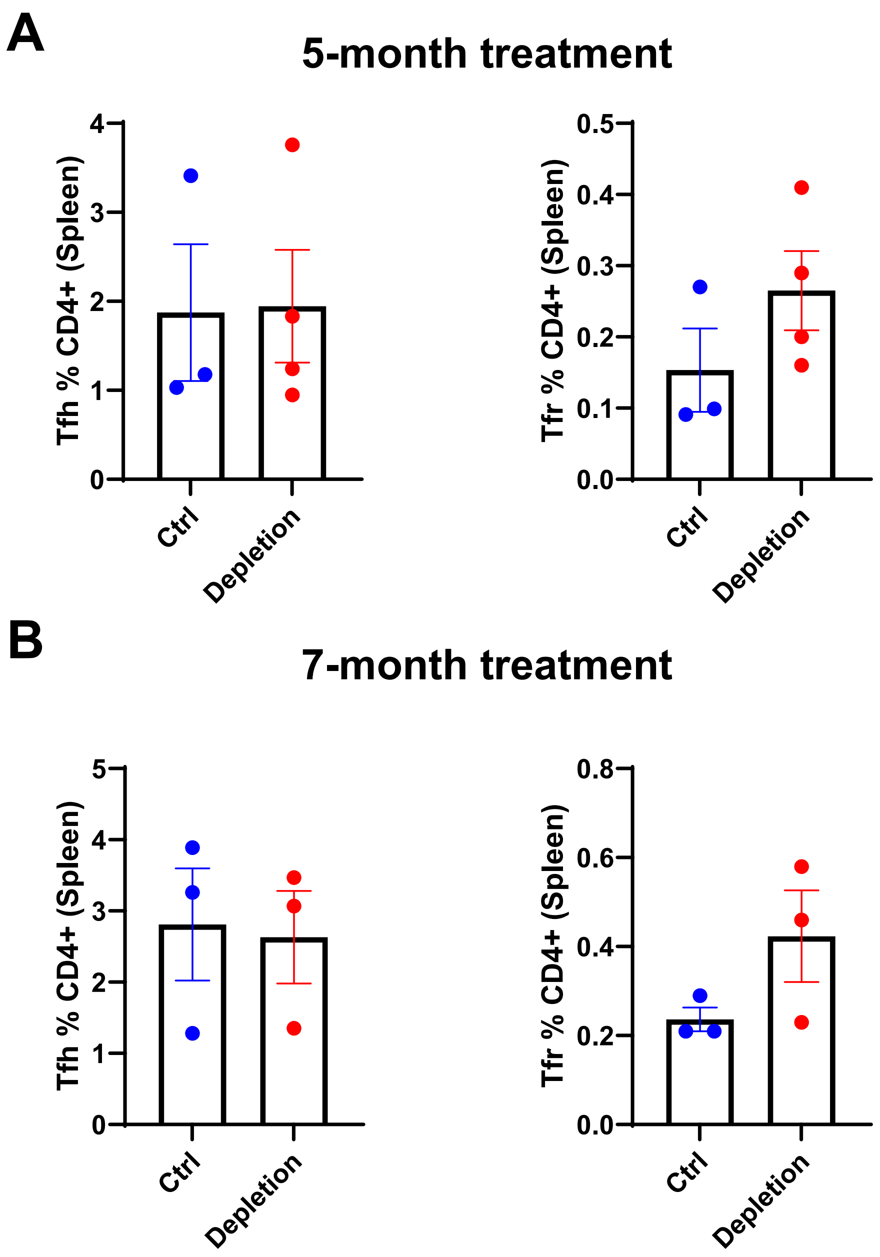


**Figure S6. Effect of *Rg* depletion on the frequency of Tfh and Tfr in splenocytes.** Frequency of Tfh and Tfr cells in splenocytes after 5- months (A) or 7 months (B) of metronidazole treatment. Mean + SEM

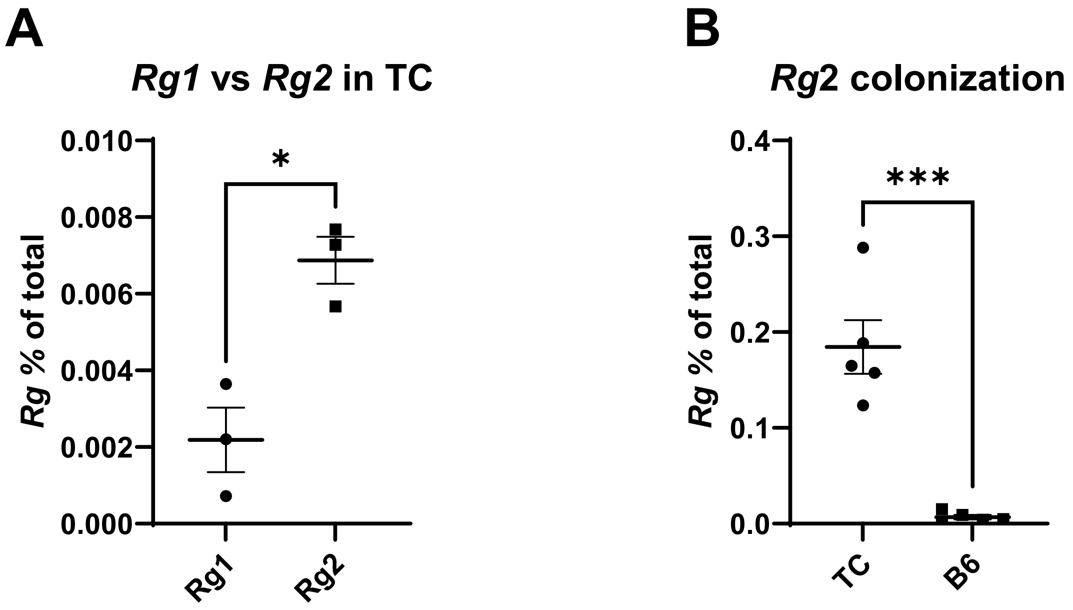


**Figure S7. TC mice were uniquely colonized by *Rg*2.** (A) *Rg*1 or *Rg*2 colonization levels in TC mice at 1 month post inoculation. (B) *Rg*2 colonization level in B6 or TC mice at 1 month post inoculation. Mean + SEM compared with t tests. *: P < 0.05; ***: P < 0.001.
